# statLens enables accurate and interpretable statistical intelligence

**DOI:** 10.64898/2026.11.16.767003

**Authors:** Dezhao Fu, Ajinkya Patil, Udbhav Kush, Ali Reza Taheriyoun, Ali Rahnavard

## Abstract

Diverse existing approaches and analyses of large-scale biomedical omics datasets along with the complexity of the corresponding statistical analytical workflows pose challenges to the direct application of large language models (LLMs) to data analysis tasks. We present statLens, a model-agnostic AI-driven statistical intelligence framework that integrates LLMs, code and human review. We demonstrate statLens using differential expression analysis across heterogeneous biomedical data as a use case. On 102 benchmark tasks from 74 datasets across six omics modalities, statLens maintained consistently high accuracy (92.5–95.0%) across seven language models and outperformed LLM-only inference (17.5–46.8%), whereas autonomous-agent accuracy varied widely (58.7–99.5%). We also showed that statLens substantially improved the accuracy of the four small language models (92.5–94.4%) over autonomous agents (58.7–92.2%) and LLM-only inference (17.5–43.5%). By decomposing analytical workflows into complementary tasks for code, LLMs and human experts, statLens provides an integrated framework for more accurate, interpretable and reproducible AI-assisted data analysis, particularly in complex omics settings.

---

The rapid expansion of statistical and computational methods in biomedical research, coupled with the growing availability of large-scale omics datasets ^1^ and rich clinical metadata, provides an opportunity to combine artificial intelligence (AI) with researcher expertise for public health and biomedical research. Rapidly evolving omics technologies, complex data characteristics, study designs, and analytical methods present challenges in deciding on the appropriate steps for data processing and conducting analyses ^2,3^, such as differential expression analysis (DEA) ^4–6^. A key unmet need is therefore a comprehensive automated platform that can standardize and coordinate statistical analysis across analytically comparable omics data types while accounting for study-specific design characteristics and data structures. Such a platform could streamline the analytical workflow and support the systematic investigation of scientifically relevant, study-specific hypotheses.

For differential analysis, key statistical decisions arise before model fitting. Analysts must select a suitable model family ^4^, specify the primary comparison ^7^, identify the experimental unit and dependencies among observations ^8^, and determine how batch variables and other covariates enter the model ^9^. These interdependent decisions must be informed by the research objective, experimental design, metadata and feature matrix. Although large language models (LLMs) can provide useful statistical reasoning ^10^ and generate code ^11,12^, supporting these decisions also involves computational tasks, such as profiling matrix values, verifying sample alignment and assessing design estimability. Statistical knowledge and code generation alone do not ensure that these computations are performed correctly or that the proposed specification addresses the intended question. These requirements motivate the development and the use of AI agents that coordinate contextual reasoning, data computation, programmatic validation and researcher review.

To date, scientific agents built on LLMs have made substantial progress in planning and executing analyses ^13,14^. AutoBA ^15^ generates analysis plans, writes and repairs code, and executes omics workflows; its evaluation includes expert review of plans, code and outputs. Robin ^16^ integrates literature agents with Finch for experimental data analysis and synthesizes multiple analysis trajectories into consensus findings. Differences between trajectories may reflect defensible choices ^17,18^, stochastic instability or statistical errors; disagreement alone does not distinguish them. Decisions encoded in generated code remain inspectable. For differential analysis, an additional goal is to make the preceding decision process explicit: which tasks require computation, which require interpretation of study context, and which decisions require researcher confirmation.

Evaluation already addresses parts of this problem ^19–21^. BLADE ^22^ assesses intermediate choices of conceptual variables, transformations and statistical models against expert-validated alternatives, and Robin evaluates its Finch agent with an RNA-seq rubric covering technical replicates, count-aware models and covariate reporting ^16^. A complementary focus is therefore the development of an evaluation strategy tailored to differential analysis of omics data. Such a strategy should assess the coherence of linked statistical choices specified before analysis and examine how those choices affect the resulting sets of significant features.

Here, we present statLens (**Fig. 1**), a model-agnostic, AI-driven statistical intelligence framework that integrates LLMs, code and human review through explicit task decomposition. statLens does not depend on a specific language model: models are interchangeable through a standard API interface, and every model uses the same prompts, rules and checks. Code computes facts from the data and validates every LLM output against implemented rules. The LLM interprets the study context and provides reasoning and recommendations, while researchers review, revise and confirm each decision. These responsibilities are coordinated through a structured end-to-end statistical analysis pipeline that combines computed data properties, confirmed study information and proposed design choices. Within the supported scope, code derives dependence labels from confirmed fields, while the LLM proposes a candidate design formula using the design card and computed data summaries. The reviewed specification is then validated and converted into an executable design for the supported analysis workflow. This decomposition connects study evidence, explicit design decisions and executable analysis within one reviewable workflow.

**Figure 1:**
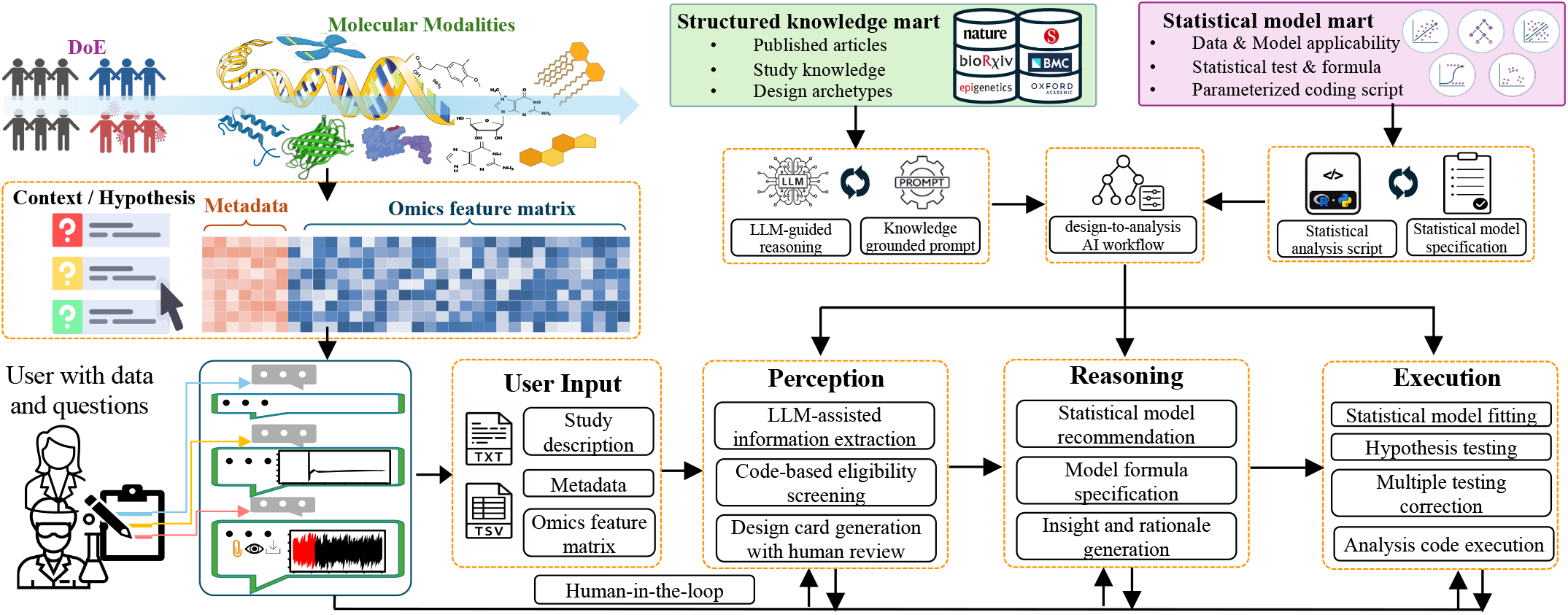
statLens enables a human-in-the-loop end-to-end workflow for differential expression analysis. Given a study description (including the scientific hypothesis), metadata, and an omics feature matrix provided by the user, statLens integrates a structured knowledge mart and a statistical model mart to support an end-to-end workflow through three modules: Perception, Reasoning, and Execution. The Perception module extracts study information, performs code-based eligibility screening, and generates a statLens design card for human review. The Reasoning module recommends an appropriate statistical model, specifies the model formula, and generates a grounded rationale. The Execution module runs the analysis code to perform model fitting, hypothesis testing, and multiple testing correction. Across all three modules, users can interact with statLens by providing comments and modifying intermediate outputs before proceeding to the next module.

To evaluate statLens from statistical design to final analysis outputs, we constructed an independent benchmark of 102 differential-analysis tasks spanning six omics modalities, following the capsule-based approach of BixBench ^21^. Source selection and prompt construction were designed to limit information leakage ^23^. To standardize scoring of otherwise open-ended design decisions, we benchmarked statLens against LLM-only and autonomous-agent baselines using five question types with predefined reference answers: data retrieval (Q1), statistical model selection (Q2), analytical objective identification (Q3), sample dependence assessment (Q4) and design formulation (Q5).

Our results showed that statLens achieved question-level accuracies of 92.5–95.0% across seven language models, exceeding LLM-only inference and showing sub-stantially less variation across models than autonomous agents. statLens particularly benefited small language models (SLMs), increasing their mean accuracy from 81.0% with autonomous agents to 93.6% and supporting accurate statistical design with locally deployable models. These findings support statLens as a practical framework for design-aware DEA and illustrate how coordinating researcher oversight, deterministic computation and constrained language reasoning can enable reliable statistical workflows.

## Results

### statLens enables end-to-end statistical analysis through collaboration among researchers, code and LLMs

Here, we define statistical intelligence as the ability to translate research questions and data into interpretable statistical results through coordinated interactions among researchers, code and LLMs. We present statLens as a real-world demonstration of this concept: an end-to-end statistical workflow in which researchers, code and LLMs have complementary roles (**Fig. 2a**). Code profiles the feature matrix, including value types, ranges, integer status and missing values; verifies sample correspondence between the matrix and metadata; and counts samples by group, time point and subject. It also executes the analysis after researcher confirmation. The LLM addresses questions that require interpretation beyond what code can determine directly. It interprets the study context, recommends study-design elements and explains the rationale for its recommendations, including why alternatives were not selected. Researchers provide the study context, review and revise recommendations, and make the final decision at each step. statLens thus organizes the three roles into a single design-to-analysis workflow. Because data computation and validation are handled by code, the language model is an interchangeable component of statLens. Researchers can select any model, from developer-hosted LLMs to locally deployed SLMs, by specifying its endpoint.

**Figure 2:**
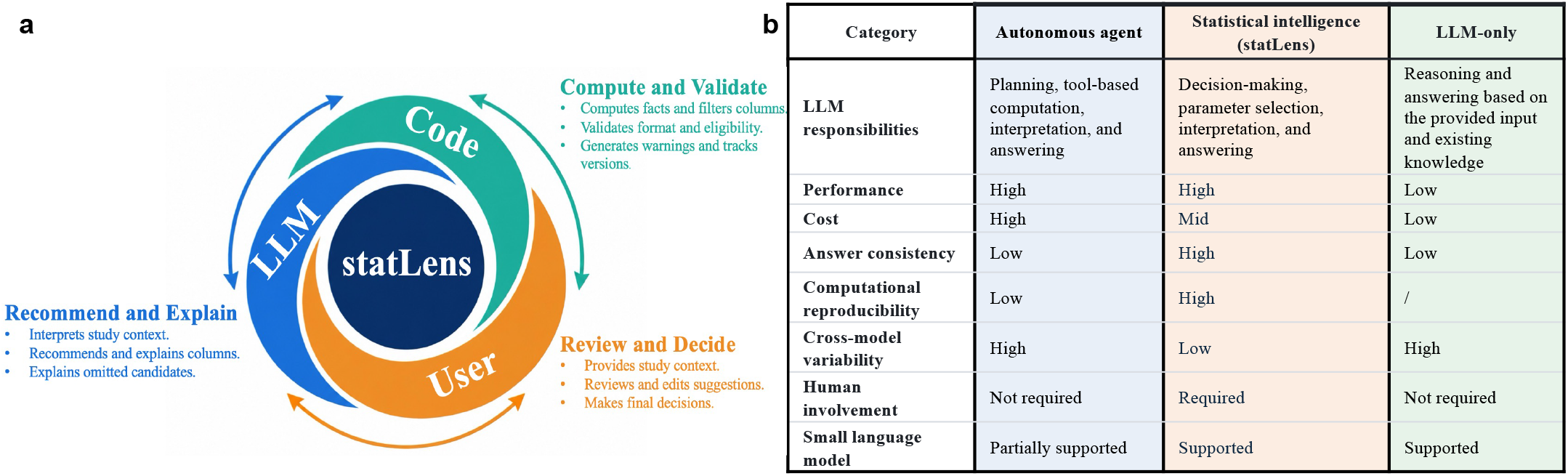
The statistical intelligence framework of statLens and comparison with alternative LLM-based approaches. **a**, Collaboration among the large language model (LLM), code, and user in statLens. The LLM interprets study context and recommends and explains column selections; code performs computations, validates inputs, and generates warnings; and the user provides study context, reviews and revises recommendations, and makes final decisions. **b**, General qualitative comparison of statLens, autonomous agents, and LLM-only approaches.

statLens coordinates interactions among researchers, code and LLMs throughout seven steps across three modules (**Fig. 1**). In the perception module (steps 1 and 2), statLens uses deterministic computations to profile the feature matrix and an LLM to extract study information, with responses grounded in evidence from the study description and checked programmatically. statLens then combines rule-based assignments with constrained LLM recommendations into a design card that defines the data context, research question and study design for researcher review and confirmation. In the reasoning module (steps 3–5), the LLM recommends a statistical model family and proposes a candidate design formula using the confirmed design card and computed data facts. Code checks the recommendation and formula against implemented consistency rules, and researchers review and confirm the input specification. The LLM then explains the reviewed decisions and their limitations, with code checking the structure and coverage of the explanation. In the execution module (steps 6 and 7), code assesses design-matrix rank, residual degrees of freedom and contrast estimability and prepares the executable design. Rule-based recommendations guide test selection, while code validates the analysis parameters. statLens then generates editable R scripts from predefined templates for model fitting, hypothesis testing and multiple-testing correction, with execution initiated by the researcher.

Two complementary resources, the statLens knowledge mart (**Supplementary Section 1.1**) and the statistical model mart (**Supplementary Section 1.2**), support the design and implementation of statLens. The statLens knowledge mart contains structured records derived from 6,855 PubMed-indexed omics articles, covering study backgrounds, experimental designs, statistical methods and reported findings. Through iterative review and refinement with LLM assistance, we used these records to develop the workflow logic, computational procedures, LLM prompts and requirements for researcher input and confirmation. The statistical model mart provides statistical specifications and reusable code implementations, including DESeq2-based analysis scripts ^24^. It brings together data requirements, model applicability conditions, model formulas and hypothesis tests with the code needed to perform the corresponding analyses. These implementations serve as parameterized templates that statLens configures using the selected model, design formula and analysis settings, producing editable scripts for model fitting, hypothesis testing and multiple-testing correction.

### statLens maintains high accuracy and answer rates across language models

To evaluate the statistical design decisions supported by statLens, we adapted the multiple-choice evaluation framework of BixBench ^21^ to a predefined benchmark comprising 102 tasks from 74 public datasets (**Supplementary Section 1.3**). The benchmark covered six omics modalities: RNA-seq (57 tasks), metabolomics (24), proteomics (7), amplicon microbiome profiling (7), small RNA (5) and DNA methylation (2). Each task provided a study description, sample metadata and a feature matrix, together with five questions assessing data retrieval (Q1), statistical model selection (Q2), analytical objective identification (Q3), sample dependence assessment (Q4) and design formulation (Q5). These 510 questions tested whether a workflow could identify the meaning and scale of the data, select an appropriate model family, characterize the primary comparison and dependence among observations, and specify a model formula and statistical test addressing the research question. We compared three paradigms supplied with the same study information (**Fig. 2b**; **Supplementary Section 4**). In the autonomous-agent condition, each model generated and executed analysis code in a Jupyter notebook ^25^, following the BixBench approach ^21^. In the LLM-only condition, models answered directly from the supplied inputs and their pretrained knowledge, without executing code. In the statLens condition, the same models provided language reasoning within the structured statLens workflow, which combined programmatic data characterization, statistical design rules and response validation (steps 1–5). Each paradigm was evaluated with seven models: three accessed through developer-hosted APIs (GPT-5.5^26^, DeepSeek-V4.1-Flash ^27^ and Claude Haiku 5.5^28^) and four small language models (SLMs; defined below): Gemma 4 26B-A4B ^29^, Qwen3.5-9B ^30^, Phi-4^31^ and Gemma 3 12B ^32^. Each model–paradigm combination was run three times independently. We report strict question-level accuracy, counting unanswered questions as incorrect, and answer rate (both defined in Section 4). Across the seven models, statLens achieved accuracies of 92.5–95.0%, compared with 58.7–99.5% for autonomous agents and 17.5–46.8% for LLM-only inference (**Fig. 3a,d–f**). The accuracy range across models was 2.5 percentage points with statLens, compared with 40.8 percentage points for autonomous agents. These results indicate that statLens maintains high accuracy with reduced sensitivity to the choice of the foundational language model.

**Figure 3:**
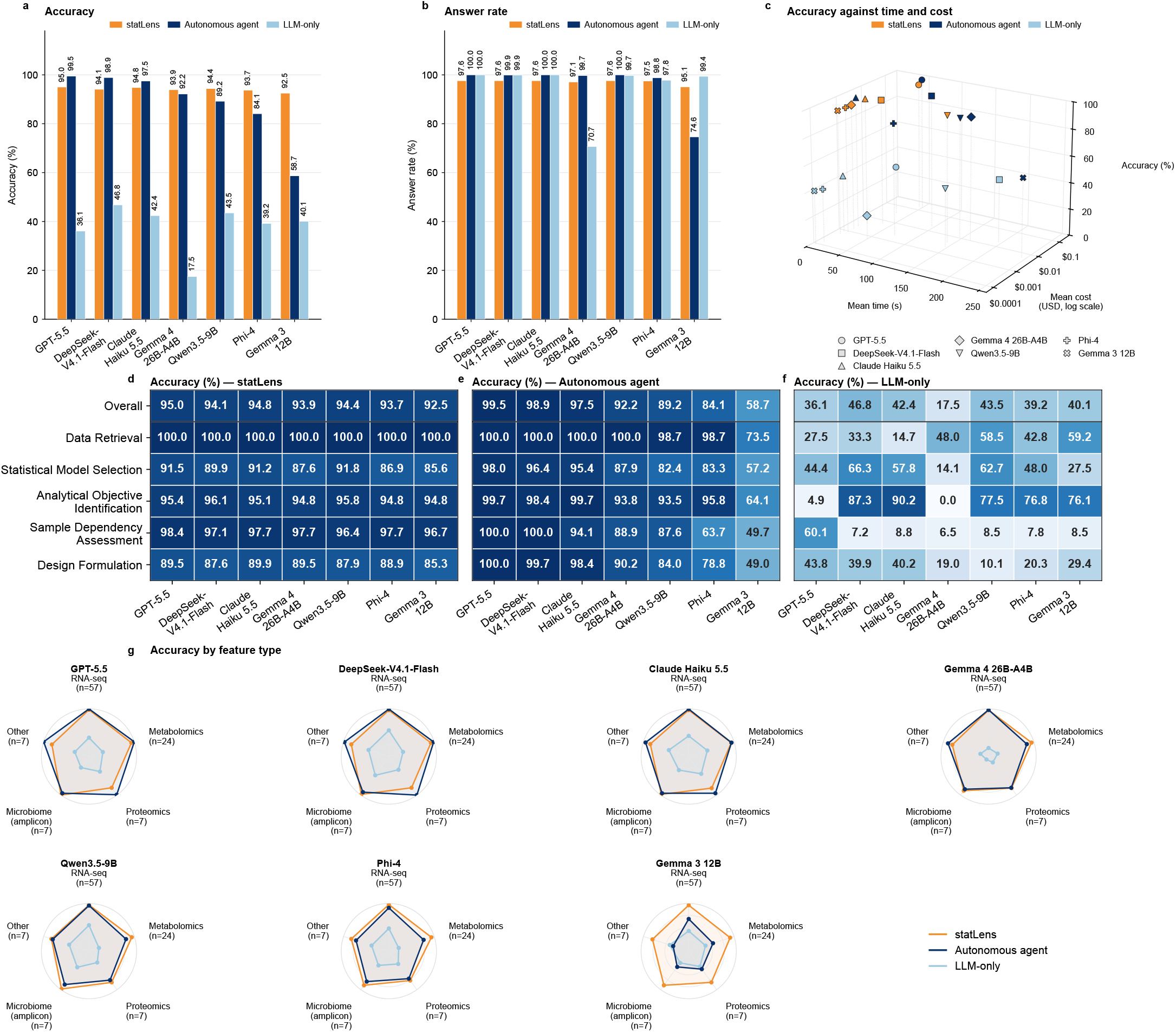
Benchmarking statLens against autonomous agents and LLM-only approaches on statistical analysis tasks. Seven language models were evaluated on 102 tasks, each comprising five multiple-choice questions. **a**, Overall accuracy, defined as the proportion of questions answered with the reference option. Missing or invalid answers and abstentions were counted as incorrect. **b**, Answer rate, defined as the proportion of questions receiving a parsable answer option, regardless of correctness. **c**, Overall accuracy in relation to mean runtime and mean cost. Each point represents a model–paradigm combination; marker shapes identify models and colors identify paradigms. Cost is shown in US dollars on a logarithmic scale. **d–f**, Overall and question-specific accuracy for statLens (d), autonomous agents (e), and LLM-only approaches (f). Cell values indicate accuracy percentages. **g**, Accuracy stratified by omics feature type for each model and paradigm. Greater radial distance indicates higher accuracy, and numbers in parentheses indicate task counts. “Other” comprises small RNA and DNA methylation tasks.

Answer rates showed a similarly narrow range across models: statLens answered 95.1–97.6% of questions, compared with 74.6–100% for autonomous agents and 70.7– 100% for LLM-only inference (**Fig. 3b**). The difference between the highest and lowest answer rates was 2.5 percentage points for statLens, compared with 25.4 percentage points for autonomous agents. This consistency extended to models with markedly different autonomousagent performance. For example, Gemma 3 12B answered 95.1% of questions with statLens, compared with 74.6% as an autonomous agent.

The remaining limitations reflected, in part, the range of designs currently supported by statLens. Twelve questions across four tasks were left unanswered because the tasks contained unmerged technical replicates, which this version of statLens does not handle. For these tasks, statLens stopped at the Design Card and requested that the technical replicates be merged before proceeding, providing no model-family recommendation, primary-contrast specification or model formula (Q2, Q3 and Q5). These exclusions set the maximum achievable answer rate at 97.6% for the current version on this benchmark.

### statLens enables accurate statistical design with small language models

SLMs are of particular interest for statistical analysis in biomedicine, but no fixed threshold separates them from large language models. A recent study considers models of 100 million to 5 billion parameters ^33^, whereas a functional definition counts any model that can run on a single consumer device with practical latency, which in 2025 meant most models below about 10 billion parameters ^34^. We therefore define SLMs operationally as open-weight models that can be served on a single graphics processing unit: dense models with at most 15 billion parameters, or mixture-of-experts models with at most 5 billion parameters active per token. Qwen3.5-9B ^30^, Gemma 3 12B ^32^ and Phi-4^31^ (14 billion parameters) are dense models within this limit, and Gemma 4 26B-A4B ^29^ activates 3.8 billion of its 26 billion parameters per token. Such models can be deployed within institutional infrastructure, allowing study descriptions, sample metadata and feature matrices to be processed without transmission to external model providers, an advantage given the privacy risks of sharing genomic data ^35^.

Without structured support, the SLMs’ performance varied substantially across models and paradigms. In the LLM-only condition, accuracy was low for all seven models (17.5–46.8%), including the four SLMs (17.5– 43.5%), and Gemma 4 26B-A4B also failed to follow the required answer format, answering only 70.7% of questions (**Fig. 3a,b**). For example, it scored 0% on analytical objective identification (Q3): although the prompt explicitly required a single-letter answer even under uncertainty, in 294 of 306 trials the model requested further details about the study instead of answering, a behavior not observed with any other model given the same prompt. As autonomous agents, the SLMs reached 58.7–92.2% accuracy, compared with 97.5–99.5% for the developer-hosted models, and Gemma 3 12B answered only 74.6% of questions (**Fig. 3a,b**). On this benchmark, direct use of the evaluated SLMs, either alone or as autonomous agents, did not consistently yield accurate statistical designs.

statLens offers a solution to this issue. Across the four SLMs, statLens increased mean accuracy from 81.0% in the autonomous-agent condition to 93.6%, a gain of 12.6 percentage points. The corresponding difference for the developer-hosted models was *−* 4.0 percentage points, yielding a difference between the two model groups of 16.6 percentage points. All four SLMs benefited from statLens: accuracy increased from 92.2% to 93.9% for Gemma 4 26B-A4B, from 89.2% to 94.4% for Qwen3.5-9B, from 84.1% to 93.7% for Phi-4 and from 58.7% to 92.5% for Gemma 3 12B (**Fig. 3a,d,e**).

Across the four SLMs, mean accuracy improved most for sample dependence assessment, followed by design formulation (**Fig. 3d–f**). With Phi-4, accuracy increased from 63.7% to 97.7% for sample dependence assessment and from 78.8% to 88.9% for design formulation. With Gemma 3 12B, the corresponding increases were from 49.7% to 96.7% and from 49.0% to 85.3%. Across all four SLMs, statLens achieved 96.4–97.7% accuracy on sample dependence assessment and 85.3–89.5% on design formulation, compared with 49.7–88.9% and 49.0–90.2%, respectively, for autonomous agents.

These gains were concentrated in decisions supported by programmatic data characterization and explicit statistical design rules. All four SLMs achieved 100% accuracy on data retrieval with statLens; for Gemma 3 12B, the corresponding autonomous-agent accuracy was 73.5%. In statLens, code computes data summaries and checks model specifications for consistency with the input data and study design, while the SLM integrates the design card, selected statistical model and computed summaries to propose candidate formulas. Researchers can then review and revise these specifications before analysis. This division of responsibilities reduces reliance on the SLM coding ability for data characterization and design validation, allowing its reasoning to focus on study interpretation and model specification. The observed accuracy gains are consistent with the benefits of supporting SLM reasoning with structured inputs, explicit design rules and programmatic checks.

The high accuracy achieved with SLMs also broadened the deployment options for statLens. Its fixed sequence of single-turn model calls used bounded prompts assembled from file summaries and returned short structured responses, constraining the model interactions required for each analysis. Autonomous-agent inference instead depended on the length of the exploration trajectory, with successive steps repeatedly supplying the conversation history and current notebook. By supporting accurate decisions with open-weight models, statLens permits deployment within institutional infrastructure, where study descriptions and metadata can remain local and model weights and inference configurations can be archived with the analysis for reproducibility. Local deployment also removes per-token API charges, while retaining the costs of hardware and computation.

### statLens reproduces androgen-associated expression patterns in mouse liver

To illustrate statLens in a real-world application, we reanalyzed GSE326073 to examine whether dihydrotestosterone (DHT) treatment altered hepatic gene expression in female mice fed a western diet ^36^, using the public release of statLens, with DeepSeek-V4.1-Flash selected as the language model. The dataset comprised eight liver RNA-seq samples collected after 8.5 weeks on the diet, with four DHT-treated and four vehicle-treated (Veh) mice. We provided statLens with the cleaned study description, sample metadata and a count matrix of 14,371 genes, for which integer raw counts were recovered from the normalized counts deposited with the published study (**Fig. 4a**).

**Figure 4:**
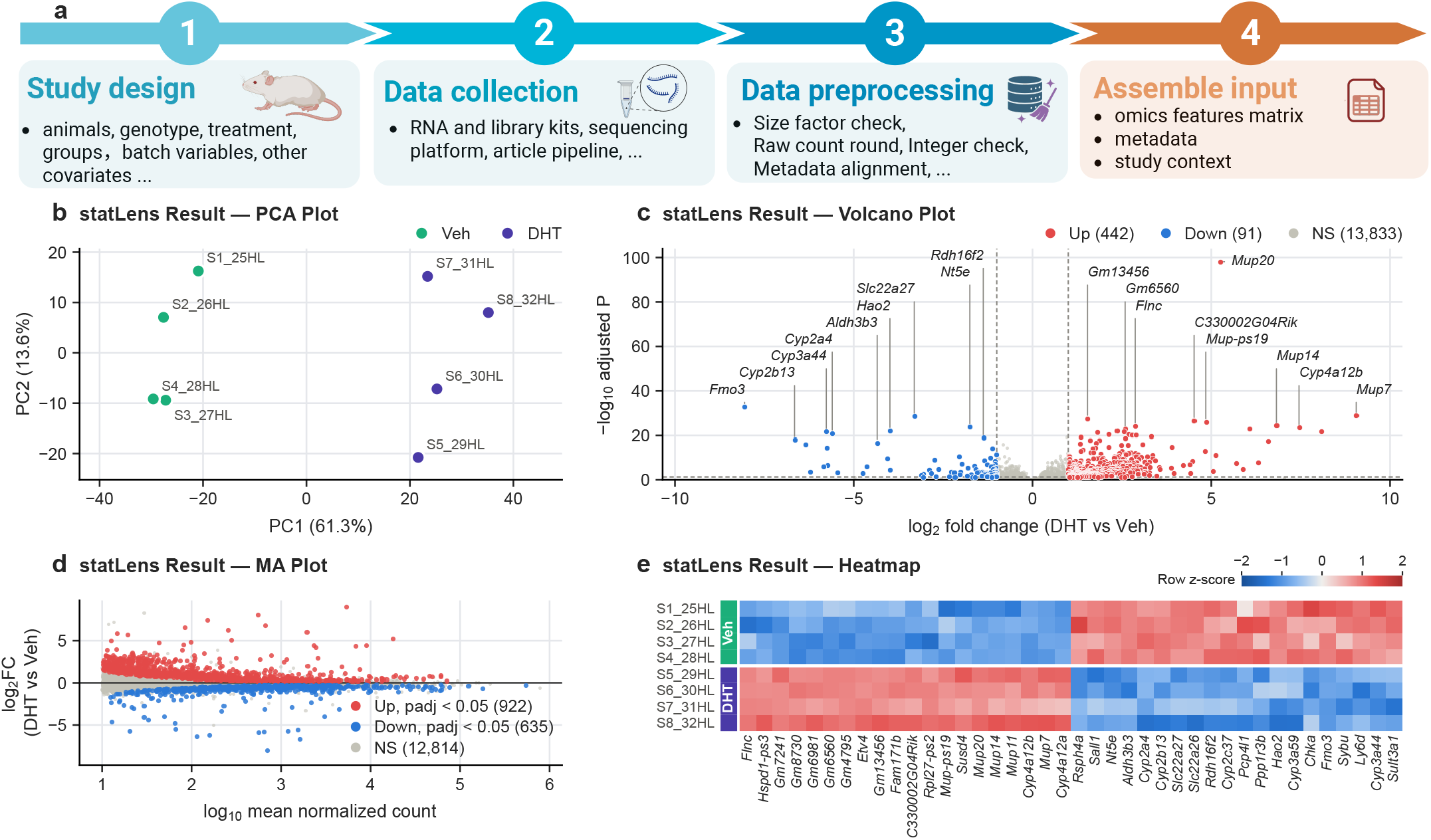
Application of statLens to a real-world RNA-seq study. Mouse liver RNA-seq data comparing dihydrotestosterone (DHT)-treated and vehicle-treated (Veh) mice, four per group. **a**, Input preparation: study-design characterization, data collection, preprocessing, and assembly of the feature matrix, sample metadata and study context. **b**, Principal component analysis of the eight samples, colored by treatment group. **c**, Volcano plot of the 14,366 genes with an adjusted *P* value. Up- and downregulated genes met both adjusted *P <* 0.05 and| log_2_ FC| *≥* 1. **d**, MA plot of all 14,371 genes, with significance defined by adjusted *P <* 0.05 alone; the five genes without an adjusted *P* value are shown as not significant. Low-count genes are absent because the deposited matrix had been filtered in the original study. **e**, Heatmap of the 20 upregulated and 20 downregulated genes in **c** with the smallest adjusted *P* values, ordered by hierarchical clustering. Colors show gene-wise *z*-scores of log_2_(normalized count + 1) across the eight samples.

We used Playwright with headless Chrome to simulate a researcher’s interactions with statLens through the entire workflow (steps 1–7), following the procedure described in Section 4. Here, the harness performed the role of the researcher in review and confirmation. Specifically, the design card (steps 1–2) identified a single-time-point comparison with one liver sample per mouse. Based on the RNA-seq context and computed count properties, the LLM recommended a negative binomial model family (step 3). Using the confirmed two-group, independent-sample design and computed data summaries, the LLM proposed the formula ∼ group (step 4). statLens code then programmatically checked its consistency with the design card. No batch variable was recorded; sex and genotype were constant, and the treatment annotation duplicated group membership, so no additional adjustment terms were included (steps 3–4). After programmatic checks and confirmation of the specification and its rationale, the DHT-versus-Veh comparison was implemented with Veh as the reference level:

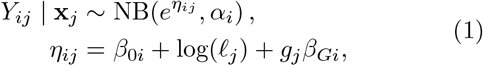

where *Y*_*ij*_ denotes the count for gene *i* in sample *j, ℓ*_*j*_ is the sample-specific normalization size factor entering as a log offset, *α*_*i*_ is the gene-specific dispersion, and **x**_*j*_ = (1, *g*_*j*_)^*⊤*^ is the design vector for sample *j*, with *g*_*j*_ = 0 for Veh and *g*_*j*_ = 1 for DHT. The coefficients *β*_0*i*_ and *β*_*Gi*_ represent the intercept and the DHT-versus-Veh effect on the natural-log expression scale, respectively. For ***β***^*i*^ = (*β*_0*i*_, *β*_*Gi*_)^*⊤*^ and *L* = (0, 1)^*⊤*^, the hypotheses were:

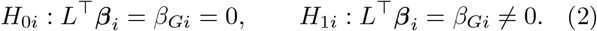

The effect size was reported as log_2_ FC_*i*_ = *β*_*Gi*_*/* ln 2.

Rule-based test recommendation identified a two-sided Wald test for this single-coefficient hypothesis (step 6). statLens generated an editable R script and executed the analysis using DESeq2 (step 7) ^24^. Here, five genes were flagged as count outliers by the default Cook’s distance check in DESeq2 and received no *P* value. Benjamini–Hochberg correction was then applied without independent filtering across the remaining 14,366 tested genes as a single multiple-testing family, with adjusted *P <* 0.05 defining significance at a 5% false discovery rate threshold ^37^.

statLens analysis result identified substantial expression differences between DHT and Veh. Principal component analysis (PCA) separated the treatment groups along the first principal component, which explained 61.3% of the variance (PC2, 13.6%; **Fig. 4b**). At adjusted *P <* 0.05, statLens identified 1,557 differentially expressed genes (922 upregulated and 635 downregulated in DHT relative to Veh; **Fig. 4d**), of which 533 also met the post-test effect-size threshold of |log_2_ FC| *≥*1 (442 upregulated and 91 downregulated; **Fig. 4c**). A heatmap of the 40 most significant of these 533 genes (20 upregulated and 20 downregulated) showed increased expression of *Mup20, Mup14* and *Cyp4a12b* and decreased expression of *Fmo3, Cyp2b13* and *Cyp3a44* in DHT-treated samples (**Fig. 4e**). These changes were consistent with the sex-biased expression of *Mup* genes, *Cyp4a12* and *Cyp3a44* in mouse liver ^38^ and the shift toward a male-biased hepatic expression profile reported in the original study ^36^.

statLens identified 1,557 differentially expressed genes at adjusted *P <* 0.05, closely matching the 1,568 reported in the original study ^36^. All 36 genes highlighted as altered in the original study, including members of the *Mup* family, were significant in the statLens analysis and changed in the reported directions. statLens used the same statistical specification as the original study: a negative binomial model with a Wald test for DHT versus Veh, Benjamini–Hochberg adjustment across all tested genes, and no batch or covariate adjustment. The small discrepancy in the number of significant genes was caused by minor numerical differences in gene-wise dispersion estimation and fitting routines between tools, and was confined to genes near the adjusted *P* = 0.05 threshold, with no effect on other analysis metrics.

## Discussion

statLens is designed as a proof-of-concept statistical intelligence framework to assist with statistical analysis. It addresses a narrow but consequential problem in omics analysis: finding an appropriate strategy for differential expression analysis (DEA) from heterogeneous study knowledge. Through this application, statLens illustrates how researchers, code and LLMs can coordinate complementary tasks to connect research questions with explicit statistical decisions and executable analyses.

The strongest benchmark gains of statLens occurred with SLMs, and its mouse liver RNA-seq application recovered published expression patterns with effect-size estimates close to an expert analysis. These results provide a proof of concept for statistical intelligence organized around a shared statistical specification. The current implementation addresses study-design interpretation, model specification and hypothesis testing, with end-to-end execution through DESeq2-based negative binomial models.

Building on advances in agent-based workflow planning and execution ^15,16^, statLens organizes collaboration around explicit representations of analytical intent and supporting evidence. Code characterizes the data and checks implemented consistency conditions; the LLM interprets study context, proposes specifications and explains recommendations; and researchers review the evidence and confirm analytical decisions. The design card connects these responsibilities by recording study information, computed data properties, the intended comparison and field provenance. Subsequent specifications make the sample subset, model formula, variable coding and hypothesis explicit. Recording the target comparison separately from reference coding helps preserve the research objective as the design is translated into an executable analysis. These intermediate representations provide a basis for tracing decisions to their supporting evidence and revisiting affected choices when upstream information changes.

The benchmark suggests that this organization can improve statistical-design accuracy, particularly for models with weaker autonomous performance. SLMs achieved a mean question-level accuracy of 93.6% within statLens, compared with 81.0% as autonomous agents. Larger developer-hosted models already performed strongly as autonomous agents and matched or exceeded statLens on the full benchmark. Nevertheless, statLens provides a guided interface for reviewing and revising analytical decisions, explicit study-specific rationales that support interpretability, and a bounded sequence of model calls with the potential to reduce inference costs. One possible explanation for the SLM gains is the allocation of data characterization and design checks to code, which reduces the computational and validation tasks delegated to the language model and allows it to focus on contextual interpretation and candidate model specifications ^39,40^. The contributions of these components remain to be established through ablation studies. Because statLens is model-agnostic, newer or domain-adapted models can be adopted as they become available without changes to the workflow. Accurate decisions with open-weight SLMs also support local deployment of statLens within institutional infrastructure. This could enable its use in settings where data cannot be transmitted to external model providers or through APIs.

The mouse liver RNA-seq application examines a complementary question: whether a reviewed specification can be carried through to interpretable results. Recovery of published expression patterns and close agreement with expert effect-size estimates support the fidelity of the configured execution path in this example. Assessing reproducibility also requires documenting the analyzed samples, hypothesis, software settings and feature-level results ^41^. Agreement in effect estimates or significantgene totals alone does not establish identical inferential conclusions. Providing an editable script alongside the design specification makes these choices available for inspection and helps researchers investigate discrepancies. The utility of this approach depends on both the models it supports and the study information it receives. Although design recommendations extend beyond the DESeq2 execution path, end-to-end execution currently remains restricted to negative binomial models. Random-effects models ^42^ and nested dependence structures are unsupported, and unmerged technical replicates prevented progression for some benchmark tasks. The work-flow also assumes sufficiently informative study descriptions, cleaned metadata and a prepared feature matrix. Missing or incorrect information about experimental units, pre-processing or potential confounding can affect subsequent decisions even when implemented consistency checks pass. These checks establish only the properties covered by their rules; yet they cannot establish the completeness of the supplied information or the scientific adequacy of every modeling assumption.

The evaluation places further boundaries on the interpretation of performance. The multiple-choice benchmark measures agreement with predefined reference specifications under controlled input conditions, and acceptable alternative analyses were not counted as reference-correct. Scope-related nonresponses also contributed to the reported errors, emphasizing the need to distinguish unsupported tasks from incorrect decisions within the supported scope. Because the automated harness advanced the workflow without substantive researcher corrections, these experiments do not quantify the additional benefit of human review, which cannot be assumed given automation bias ^43^ and evidence that human–AI combinations on average perform worse than the better of humans or AI alone ^44^. User studies are needed to assess researcher interventions, and broader execution benchmarks are required to determine how design-decision accuracy translates into agreement in downstream statistical results.

## Online methods

### statLens development

We developed statLens in three stages: resource assembly, information organization and synthesis, and iterative implementation. We first assembled two complementary resources: the statLens knowledge mart (**Supplementary Section 1.1**) and the statistical model mart (**Supplementary Section 1.2**). The knowledge mart is primarily article-based and contained structured records derived from 6,855 PubMed-indexed omics articles, covering study backgrounds, experimental designs, statistical methods and reported findings. The statistical model mart is primarily code-based and combined model fitting requirements, model applicability conditions, model formulas and hypothesis tests with reusable code implementations.

With assistance from Claude Sonnet 5.5, we organized the literature-derived information into JSON records describing data contexts, research questions, study designs and statistical analyses. This model was used only as a curation aid during development and is not part of the statLens workflow. For the statistical model mart, we summarized the statistical specifications and implementation requirements. We used these materials to identify the analytical decisions statLens would support, the information required for each decision and the respective responsibilities of code, LLMs and researchers. This synthesis informed the workflow logic, computational procedures, LLM prompts and requirements for researcher input and confirmation.

We iteratively developed and reviewed the workflow, code and prompts using the study contexts, experimental designs and statistical methods recorded in the knowledge mart. The statistical model mart supplied specifications and reusable code for implementation. The resulting workflow comprised seven steps across three modules: Perception (steps 1–2), Reasoning (steps 3–5) and Execution (steps 6–7). Code profiled the input data, checked LLM responses against implemented rules, derived dependence labels and generated and checked editable model formulas from researcher-confirmed design card fields. It also provided rule-based test recommendations and generated editable analysis scripts from predefined templates. LLM prompts supported extraction of study information with supporting evidence, recommendations for study-design elements and statistical model families, and structured explanations of decisions and their limitations. Researchers reviewed, revised and confirmed decisions at the relevant steps and initiated script execution.

In the current software release, statLens supports statistical design specification (steps 1–5) for four primary comparison structures: group-by-time interaction (GT), time course (TC), multi-group comparisons involving three or more groups (MG), and basic two-group comparisons (BG). Other comparison structures are classified as out of scope (OOS). Subject dependence is classified as independent (IND), paired or repeated (PR), or OOS. IND designs omit subject terms, whereas PR designs account for subject effects through fixed effects. Random-effects models, mixed-effects models and nested dependence structures are out of scope (OOS). Modelfamily recommendations cover negative binomial generalized linear models (GLMs), Gaussian linear models, Poisson GLMs and beta regression, with unsupported cases classified as OOS.

End-to-end execution is currently restricted to DESeq2-based negative binomial models, using Wald tests or likelihood-ratio tests (LRTs) as appropriate to the specified hypothesis. Design-matrix rank and estimability checks, model fitting, hypothesis testing and multiple-testing correction are implemented for this DESeq2 execution path.

### statLens Design Card

We developed the statLens Design Card as a standardized, machine-readable JSON representation of design-relevant information from the study context, sample metadata and omics feature matrix. As the output of the Perception module (steps 1–2), it organizes supplied study information and computed data properties, identifies unresolved design fields and provides a structured basis for researcher review and subsequent statistical design specification.

As detailed in **Supplementary Section 2**, the design card comprised 47 design fields and a separate field_provenance object recording the provenance of each field. The fields were organized into four domains: study context, assay characteristics, matrix organization and dependence, and the target comparison. Together, these domains captured the measurement properties, experimental units, dependence structure and scientific comparison needed to support subsequent statistical analysis.

Each design field was assigned one of three provenance labels in field_provenance: sourced, for values explicitly reported or supplied in the input sources, including explicitly stated absence; derived, for values obtained from available evidence through code-based calculations, rule-based inference or evidence-supported LLM interpretation, without introducing unsupported design assumptions; or pending-user-input, for required values that were neither directly available nor derivable from the supplied evidence.

Field completion followed a prespecified sequence. Directly sourced information was used when available; otherwise, values were derived from supporting evidence. Required fields that remained unresolved were marked as pending-user-input for researcher completion. Researchers reviewed, revised and confirmed the design card before proceeding to the Reasoning module.

### statLens architecture design

statLens implements a seven-step, LangGraph-based workflow across the Perception, Reasoning and Execution modules. It accepts a free-text study description, a sample metadata table and an omics feature matrix. Each module produces a specification or output used by the next: a confirmed design card, a reviewed statistical specification, and an executable analysis with results. Changes to upstream information require the affected computations, specifications and explanations to be updated before execution.

### User input preparation

Researchers prepare inputs within the supported data, study-design and analysis scope described above. The metadata and feature matrix are uploaded as tab-separated .tsv or .txt files with matching sample identifiers. The default upload size limit is 100 MB.

The metadata should identify the comparison variables and, where applicable, subjects, experimental blocks, batches and other covariates. Researchers specify the measurement type and current scale of the feature matrix, such as raw counts, normalized abundances, log-transformed values, intensities or relative abundances, and document preprocessing already performed. The study description states the biological setting, experimental design, analytical objective and hypotheses or research questions. statLens is model-agnostic. The language model is configured in the application settings by its name, endpoint URL and API credentials, and any model served through a standard API endpoint, whether developer-hosted or deployed locally, can be used. All models use the same prompts, rules, validation checks and workflow.

### Perception module

The Perception module (steps 1–2) combines computational data profiling with contextual information extraction to construct the design card.

#### Step 1: Information collection

Code profiles the complete uploaded feature matrix, recording value types, ranges, integer status and missing values. It checks sample correspondence between the matrix and metadata and summarizes available group, time-point and subject annotations. The LLM receives the study description and computed file summaries and extracts design-relevant information with supporting evidence from the description. Code checks the returned information against implemented response and consistency rules before it is used to populate the design card.

#### Step 2: statLens Design Card generation

Computed data facts and extracted study information are combined to populate the design card. Predefined rules assign design labels, including the primary-contrast class and subject-dependence category, while the LLM proposes values for fields requiring contextual interpretation. Field values and provenance follow Section 4; unresolved required fields are marked pending-user-input for researcher clarification.

The module applies the supported-scope rules and flags requirements that must be addressed before progression, including the need to merge technical replicates. Researchers review the draft card, correct or supplement its information and confirm the intended comparison. Provenance is updated following these revisions, with explicitly supplied researcher information labeled sourced. The confirmed card is passed to the Reasoning module.

### Reasoning module

The Reasoning module (steps 3–5) converts the confirmed design card into a reviewed statistical specification comprising the model family, analyzed sample subset, design formula, variable coding and primary comparison. Design checks inform both formula revision and the explanation of the final specification.

#### Step 3: Statistical model recommendation

Code computes summaries relevant to model assessment for the samples entering the analysis, including zero prevalence, sample-level totals and mean–variance relationships within relevant groups. The LLM evaluates the supported model families using these summaries together with the assay, measurement scale and preprocessing information in the design card. Numeric properties are interpreted in the context of the measurement process and study information.

The structured recommendation identifies a model family, modeling scale and any required transformation, and explains the suitability or limitations of the candidate families. Unsupported cases are classified as OOS. Code checks consistency with recorded data facts and implemented workflow constraints. Researchers select and confirm the model family.

#### Step 4: Model specification

For the analyzed sample subset, code checks sample correspondence between the feature matrix and meta-data, summarizes metadata variables and missing values, and tabulates the observed group–time–subject structure. The LLM uses these findings, the confirmed design card and the selected model family to propose a candidate design formula. The formula represents the primary comparison and, where applicable, subject or blocking effects, batch variables and other covariates.

Code checks the candidate formula against implemented syntax and design-consistency rules, including variable availability and agreement with the confirmed design fields. The displayed specification identifies the role of each term, variable coding and reference levels. The primary comparison is recorded separately from the choice of reference coding.

Researchers can edit the formula or revise the underlying design card, after which the affected checks are repeated. They confirm the input formula, analyzed sample subset, coding and primary comparison before proceeding. Design-matrix rank, residual degrees of freedom and contrast estimability are assessed in step 6.

#### Step 5: statLens Insight

The LLM generates statLens Insight from the confirmed design card, computed data facts, selected model family and reviewed input specification using a predefined structured prompt. It explains the primary comparison, subject dependence, batch effects, other covariates and design formula. For each decision, it states the supporting study-specific evidence, assumptions and remaining uncertainties, including the implications of including or omitting adjustment variables.

Code checks the response structure and coverage of the required topics. Researchers review the explanation alongside the input specification and can request revisions or correct design information. Changes to the specification require the affected checks and explanations to be updated. The reviewed input specification and its rationale are passed to the Execution module.

### Execution module

The Execution module (steps 6–7) validates and converts the reviewed input specification into an executable design, analysis settings and an R script. Model fitting and hypothesis testing occur when the researcher initiates execution in step 7. The current public release supports DESeq2-based negative binomial models; workflows requiring other execution implementations stop after the Reasoning module.

#### Step 6: Model fitting

This step prepares the executable design and configures model fitting and hypothesis testing. Code constructs a design matrix *X* from the reviewed input formula and metadata for the analyzed samples using R formula rules. It checks matrix rank, residual degrees of freedom and whether the intended primary comparison is represented and estimable under the specification. Residual degrees of freedom are calculated as

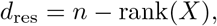

where *n* is the number of analyzed samples.

Where the input formula contains redundant terms, predefined rules generate an executable parameterization by removing redundant representations and introducing explicit derived columns where required. These transformations preserve the modeled mean structure and the estimable primary comparison. The executable formula and final matrix are rechecked following any transformation. If the intended comparison remains non-estimable or the design otherwise fails validation, researchers are directed to revise the relevant design decisions before proceeding.

The interface displays the input and executable formulas, mappings to the count matrix and metadata, and any excluded variables or added columns with their explanations. Researchers review these changes alongside statLens Insight. If a change affects the explanation of the specification, the affected text is revised before final confirmation.

Code uses predefined rules to recommend a statistical test based on the confirmed primary comparison and executable design. For DESeq2, the available tests are the Wald test and likelihood-ratio test (LRT). A Wald test uses an explicit contrast expressed in the executable coding. An LRT uses full and reduced models whose difference represents the intended hypothesis. Code validates test-specific requirements, including that the reduced model is nested within the full model for the same analyzed samples.

Researchers select the test and configure the size-factor estimation method, dispersion-trend fitting method and result-extraction parameters. The displayed defaults for sfType and fitType are ratio and parametric, respectively. The interface distinguishes design-derived settings from software defaults. Code checks parameter values and consistency with the executable specification. Changes affecting the sample subset, formula, coding or primary comparison require the affected design checks and explanations to be updated. Researchers confirm the executable design, test specification, analysis parameters and validation findings before proceeding.

The default statLens configuration applies Benjamini– Hochberg (BH) correction at a false discovery rate thresh-old of 5%, with independentFiltering = FALSE. For each primary comparison, all tested molecular features constitute one multiple-testing family. Under these settings, features with valid BH-adjusted *P <* 0.05 are considered statistically significant ^37^.

#### Step 7: Analysis code generation and execution

Code configures predefined R templates from the statistical model library using the submitted data, confirmed executable design and validated analysis settings. The script implements sample alignment and subsetting, size-factor and dispersion estimation, negative binomial model fitting, the specified Wald test or LRT, multiple-testing correction, significant-feature selection and result visualization ^24^.

Researchers can inspect, edit, copy or download the script before running it. When a run is initiated, the software executes the script currently displayed in the editor. Researchers review analysis-relevant edits before execution and can revise and rerun the script.

The module returns differential-feature tables containing effect estimates, nominal and adjusted *P* values, significant-feature tables, normalized count matrices and analysis summaries. Visualizations depend on the analysis and available results and may include volcano plots, MA plots, principal component analysis plots, heatmaps and top-feature summaries.

### Evaluation design

#### Design-decision benchmark

We constructed a benchmark of 102 statistical-design tasks from 74 public datasets to evaluate how models specify an omics analysis from study context and data. The benchmark includes diverse omics tasks from RNA-seq, metabolomics, proteomics, amplicon microbiome, small-RNA and DNA-methylation studies. We adapted the analysis-capsule approach used in BixBench ^21^. Benchmark construction followed three stages: article retrieval and screening, capsule construction, and question generation and validation.

#### Article retrieval and screening

We screened published studies and their public data records for accessible omics matrices, sample meta-data and sufficient analytical context. Data sources included the Gene Expression Omnibus (GEO) ^45^, Array-Express ^46^, PRIDE ^47^ and MetaboLights ^48^. Source publications provided the research questions and experimental context, while repository records supplied the data and sample annotations. Each task defined an analysis question and sample scope within a dataset. To avoid data leakage, we also excluded datasets from studies included in the statLens knowledge mart.

#### Capsule construction

Following the analysis-capsule approach of BixBench, an expert constructed a capsule for each of the 102 tasks with assistance from Claude Sonnet 5.5, using the articles and repository records identified during screening. Each capsule contained a cleaned omics feature matrix, corresponding sample metadata, a study description and a reference Jupyter notebook. The notebook documented data preparation and the task-specific analysis workflow, including executable code, recorded outputs and inter-pretations relevant to the research question. The expert checked the data and sample annotations against the source records and reviewed the notebook to establish reference material for subsequent question generation and validation.

**Figure 5:**
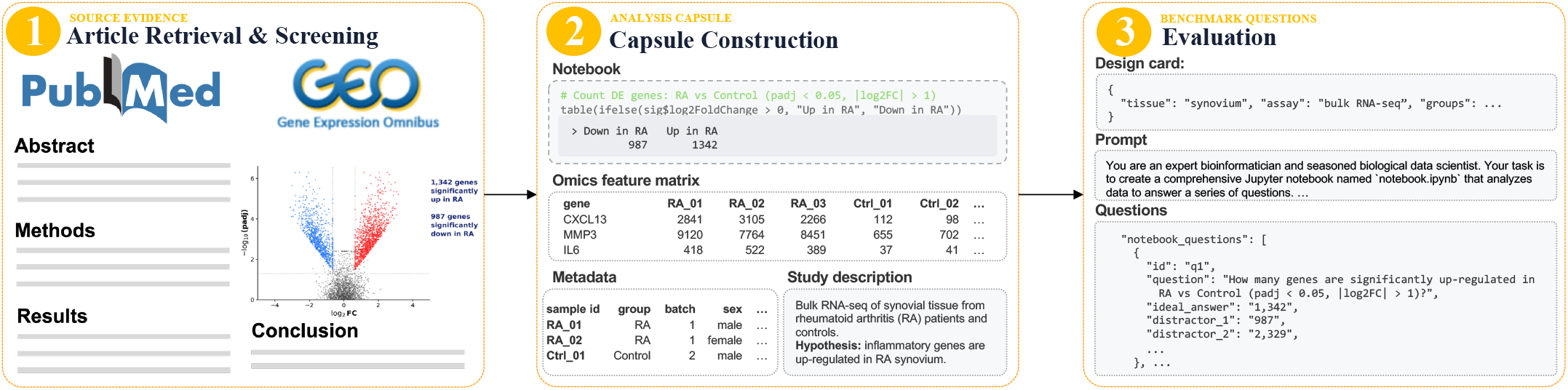
Construction of the statistical design-decision benchmark. The benchmark was developed through three stages: article retrieval and screening, analysis capsule construction, and evaluation question generation. Each capsule comprised a study description, sample metadata, an omics feature matrix and a reference analysis notebook. A reference design card was used to construct five multiple-choice questions assessing data retrieval, statistical model selection, analytical objective identification, sample dependence and design formulation.

The expert formulated research questions supported by the available measurements, sample structure and biological context. These questions could differ from those investigated in the source publications. A dataset could therefore contribute multiple tasks addressing different comparisons, groups, time points or sample subsets. Study descriptions followed a common writing guide and used the investigator’s voice. Each description stated the task-specific analytical objective, intended comparison, reference group and sample scope, together with relevant information about the biological context, assay, matrix values and relationships among samples. Here, we did not disclose information related to model formulas or statistical test labels in the study descriptions to avoid data leakage.

#### Question generation and validation

An expert constructed a reference design card containing 47 design fields for each task. Each card was checked for consistency with the task’s cleaned omics feature matrix, sample metadata, study description and reference Jupyter notebook. The reference cards were then used to construct five multiple-choice questions per task, yielding 510 questions across the 102 tasks.

Q1 assessed the representation of matrix values using five substantive categories: unnormalized counts, library-size-normalized values, continuous values on a linear scale, continuous values on a logarithmic scale, and proportions strictly between 0 and 1. Q2 assessed model family and modeling scale through six alternatives: a negative-binomial GLM for counts, a Poisson GLM, a Gaussian model after a log-like transformation, a Gaussian model on the original scale, beta regression, or another approach, such as a censored model, a zero-inflated model or a rank-based method.

Q3 assessed the primary-comparison structure: a basic two-group comparison, a multi-group comparison involving at least three groups, a time-course comparison, a group-by-time interaction, or none of these structures. Under the prespecified convention, a comparison between two specified levels was classified as a two-group comparison. Q4 assessed sample dependence and the unit of analysis. Alternatives represented independent observations, two task-specific fixed-effect specifications using designated metadata columns to identify paired or repeated units, and aggregation of nested units or random-effects modeling. Q5 assessed the design formula and its associated hypothesis test.

For Q5, the four substantive alternatives comprised the reference specification, a specification containing error *D*_1_, one containing error *D*_2_, and one containing both errors. Each error was selected from a predefined catalogue of individual perturbations: omission or unjustified inclusion of unit effects; pairing by replicate number; omission of relevant batch or covariate adjustment; an incorrect target comparison or reference group; a mismatch between pairwise and omnibus testing; inappropriate use of additive or interaction terms; inclusion of a constant metadata column; or construction of a saturated model. At most one of the two errors could alter the hypothesis-test specification.

Answer options were annotated as reference-correct, acceptable alternatives, suboptimal, incorrect or abstention. Accuracy counted only reference-correct answers.

Incorrect options carried tags identifying their error types. Every question included an abstention option, which was always scored as incorrect. Substantive options were shuffled using reproducible seeds derived from the task, question and evaluation-replicate identifiers. Q5 alternatives used neutral identifiers, and the abstention option was always placed last.

Claude-based review agents examined the question bank with access to the complete task capsules, reference design cards, source publications and the Internet. Their selected answers were compared with the reference answer keys. Agreement with the reference key was recorded for 510 of the 510 questions.

Finally, a separate overlap audit compared the benchmark source studies and datasets with those represented in the statLens knowledge mart. No overlapping studies or datasets were identified.

### Agent evaluation

The agent, LLM-only and statLens conditions received the same three inputs for each task: the study description, sample metadata and omics feature matrix. Across all conditions, questions left unanswered because the model’s input-token limit was exceeded were retained in the evaluation and scored as incorrect.

We adapted the BixBench evaluation environment and open-source agent framework ^21,49^. The system prompt was based on its multiple-choice capsule template, modified for sequential questions about statistical design for differential analysis. The task prompt contained the study description, workflow instructions, required answer format and current question, and explicitly prohibited Internet searches.

Agents had access to three tools: edit_cell, list_workdir and submit_answer. After each edit, the entire Jupyter notebook was re-executed from the first cell in a Python kernel, with R accessed through rpy2. File access was restricted to the current run directory to prevent access to other evaluation materials.

Questions were presented sequentially. Unparsable submissions could be retried up to twice; a submission that remained unparsable after these retries was recorded as invalid. Once a question was finalized by a parsable answer or an invalid result, it was locked before the next question was presented. Each agent run ended when all five questions had been finalized, 30 model calls had been made, or 90 minutes had elapsed, whichever occurred first.

GPT-5.5, DeepSeek-V4.1-Flash and Claude Haiku 5.5 used provider-default reasoning settings and automatic tool selection. Claude Haiku 5.5 was accessed through its native API at the default temperature, with explicit instructions to use tools within the notebook workflow. Qwen3.5-9B, Gemma 3 12B, Phi-4 and Gemma 4 26B-A4B were accessed through OpenRouter with tool calling required.

Each model–task combination was run independently three times, with substantive answer options reshuffled for each replicate. Reruns were restricted to infrastructure failures unrelated to input-token limits. All other runs were included in scoring, including runs that answered only some questions or submitted no answers.

### LLM-only evaluation

In the LLM-only condition, models could directly inspect the supplied study description, metadata and feature matrix. Notebook access, tool calling and code execution were disabled. Each question was submitted separately, using the same option order as the corresponding agent run.

The prompt required models to read the complete study description and inspect metadata and feature-matrix content selectively according to the information needed for each question. Answers relied on the model’s prior statistical knowledge and information read directly from these inputs.

Models returned their selected options directly in the required format. Answer parsing, resubmission and scoring followed the agent evaluation protocol. Model versions, providers, reasoning settings and temperature settings also matched the agent condition.

### statLens evaluation harness

We used Playwright with headless Chrome to simulate a researcher’s interactions with statLens. Each episode started with a fresh application instance. The harness uploaded the feature matrix and sample meta-data, entered the study description, tested the model connection and selected Extract design.

The harness accepted answers already populated by the model and selected Suggest only for fields that remained empty. Failed model calls were retried up to two additional times. Model versions, providers and applicable reasoning and temperature settings matched those used in the agent evaluation.

For the design-decision benchmark, the harness ran statLens through the first five workflow steps. Prespecified rules mapped the application outputs to the five benchmark answer options. The mapped answers followed the same answer format and scoring rules as the agent condition. A question was scored as incorrect if the application did not reach the relevant stage or its output could not be mapped to a valid option.

Only model calls associated with answering the five benchmark questions were included in the reported token usage and cost totals. This accounting scope focused the comparison on the decisions assessed by the benchmark.

### Evaluation analysis

For the multiple-choice benchmark, accuracy measured the proportion of questions answered correctly. Let *N* denote the total number of evaluated questions, *y*_*i*_ the gold-standard option for question *i*, and *ŷ*_*i*_ the parsed model response. Unanswered questions, invalid responses and explicit abstentions were represented by *ŷ*_*i*_ = ∅ and counted as incorrect. We calculated accuracy as

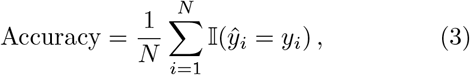

where I(*·* ) denotes the indicator function.

Answer rate measured the proportion of questions for which the model returned a parsable answer option, regardless of correctness:

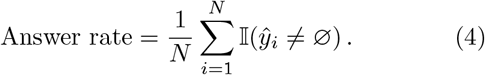

Model costs were calculated from per-call token usage and the applicable prices. Most GPT-5.5 calls used the provider’s discounted Flex service tier and were priced accordingly. Standard list prices were checked on 19 October 2026. For models accessed through OpenRouter, we used the actual call costs reported by OpenRouter whenever available.

## Declarations

### Data availability

All datasets analyzed in this study are publicly available from GEO, ArrayExpress, PRIDE and Metabo-Lights. Accession numbers for the 74 source datasets are listed in **Supplementary Section 1**. The mouse liver RNA-seq data are available from GEO (accession GSE326073). The design-decision benchmark is available at HuggingFace https://huggingface.co/datasets/omicseye/statLens-evaluation-capsule. It includes the 102 task capsules (study descriptions, sample meta-data, feature matrices and reference notebooks), reference design cards, answer keys and per-run model responses.

### Code availability

statLens is made available through a web portal at https://statlens.ai. The evaluation harness, scoring scripts, and figure-generation code are available at https://github.com/omicsEye/statLens.

For every analysis, statLens generates a standalone, editable R script based on DESeq2, so results can be reproduced and audited without the portal. The work-flow logic, decision rules, prompt templates, and analysis templates are described in **Supplementary Sections [1–4]**.

The statLens source code is subject to a pending patent application. It will be shared confidentially with editors and reviewers during peer review, and is available to academic researchers from the corresponding author on request under a non-commercial license agreement.

### Author contributions

A.R. conceived the study. D.F., A.R.T., and A.R. implemented the approach and performed performance evaluations. D.F. and U.K. tested and benchmarked various small and large language models within statLens. A.R. and A.R.T. guided data interpretation and structured the results. A.P. and A.R. designed and implemented a portal with an interface and computing infrastructure to access the models. All authors drafted, reviewed, discussed, and approved the final manuscript.

