## Supplemental Information for "statLens enables accurate and interpretable statistical intelligence"

### Supplementary materials for: statLens enables reliable and interpretable statistical intelligence

[1]Computational Biology Institute, Department of Biostatistics and Bioinformatics, The George Washington University, Washington, DC, 20052, USA.
[2]seqSight, Washington, DC, 20011, USA.

Contributing authors:;

### Section 1. Data Collection

#### Section 1.1 statLens knowledge mart

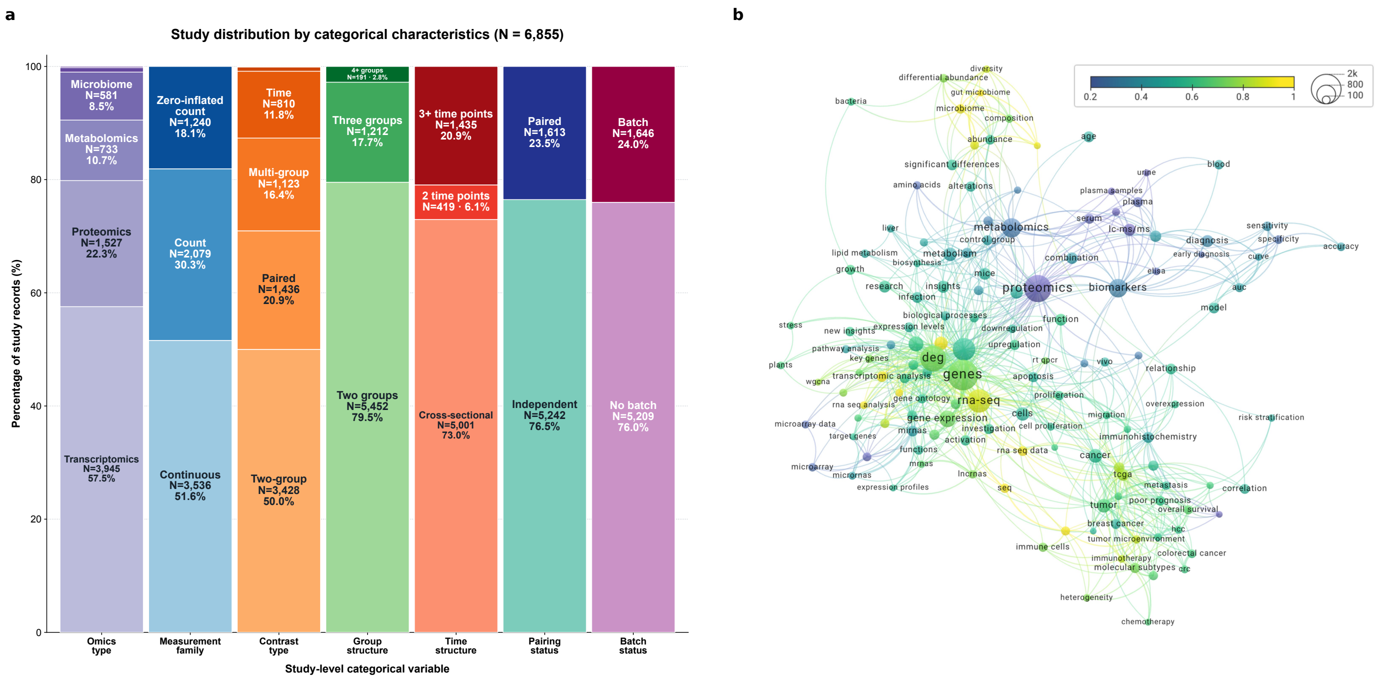


**Supplementary Figure 1.1. Composition and biomedical key-word network of the statLens Knowledge Mart. a,** Distribution of the 6,855 curated records across seven categorical characteristics. Bar segments indicate the percentage of records in each category; labels report category names, counts, and percentages. **b,** Co-occurrence network of biomedical key words extracted from the source articles. Node size represents term frequency, edges indicate within-study co-occurrence, and node color represents the term relevance score.

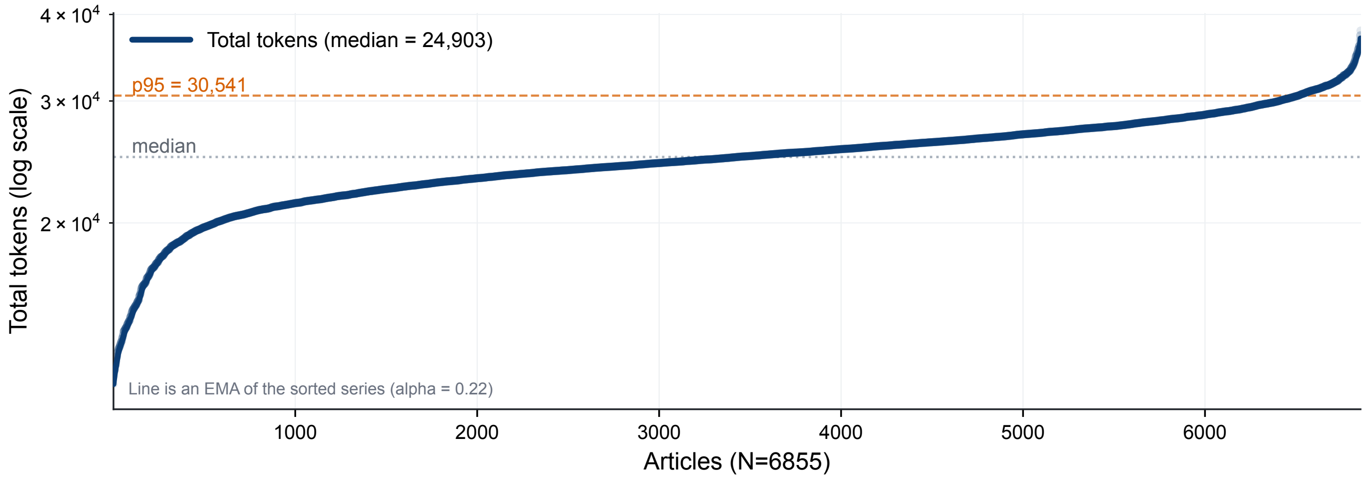


**Supplementary Figure 1.2. Overview of the token usage for statLens Knowledge Mart construction.**

**Supplementary Table 1.1.** Distribution of the 6,855 studies with a Design Card across six omics categories.

| Omics category | Representative modalities or assay types | N studies | % of corpus |
|---|---|---|---|
| Genomics | SNP genotyping and association arrays, copy-number (array CGH/SNP array) profiling, whole-genome/exome sequencing, and DNA methylation or chromatin assays | 54 | 0.79% |
| Microbiome | 16S rRNA amplicon sequencing (ASV/OTU profiling), shotgun metagenomics, and ITS amplicon profiling | 581 | 8.48% |
| Proteomics | TMT/iTRAQ, and DIA LC-MS/MS; affinity platforms (Olink, SomaScan, RPPA); and 2-DE/MALDI-TOF profiling | 1,527 | 22.28% |
| Metabolomics | Untargeted and targeted LC-MS, GC-MS, and NMR metabolite profiling, and lipidomics | 733 | 10.69% |
| Transcriptomics | Bulk RNA sequencing, expression microarrays, single-cell/single-nucleus RNA sequencing, miRNA profiling, and spatial transcriptomics | 3,945 | 57.55% |
| Other | Glycomics, DNA methylation profiling, functional metagenome profiling, and image-based cell profiling | 15 | 0.22% |
| Total | – | 6,855 | 100.00% |

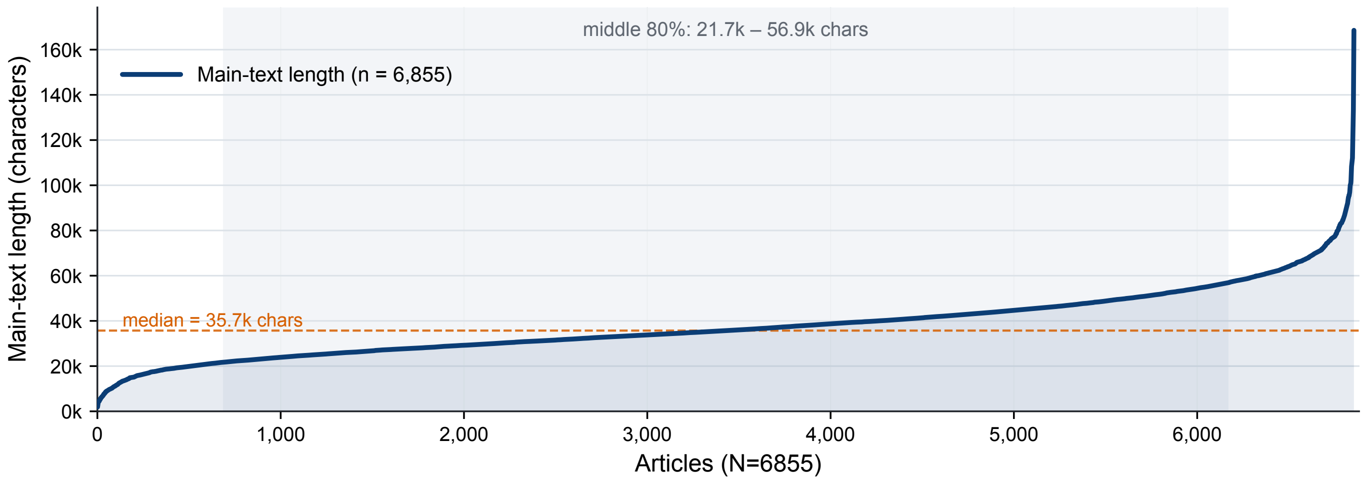


**Supplementary Figure 1.3. Overview of the text length for statLens Knowledge Mart construction.**

#### Section 1.2 statistical model mart

**Supplementary Table 1.2** Support for the statistical configurations considered by statLens in statistical model mart

| Method | Statistical model | Subject dependence | | Batch / other covariates | Hypothesis test | Multiple-testing correction[c] |
|---|---|---|---|---|---|---|
| | | Independent | Subject fixed effects | | | |
| DESeq2 [a] | NB GLM | Yes | Yes | Yes | Wald test; LRT | BH |
| NBPSeq | NB GLM | Yes | Yes | Yes | Wald test; LRT | BH |
| TSPM | Poisson GLM[b] | Yes | Yes | Yes | Poisson LRT; quasi-likelihood-adjusted test | BH |
| edgeR | NB GLM | Yes | Yes | Yes | LRT; quasi-likelihood F-test | BH |
| limma | Gaussian Linear Model | Yes | Yes | Yes | Moderated t-test; moderated F-test | BH |
| dream | Gaussian Linear Model | Yes | Yes | Yes | Moderated t-test; moderated F-test | BH |

*Notes.* NB, negative binomial; GLM, generalized linear model; LRT, likelihood-ratio test; BH, Benjamini–Hochberg correction. Entries indicate supported configurations rather than software defaults. Subject, batch, and other covariates are represented as fixed effects in an identifiable design matrix.

[a] The execution module of statLens (model fitting, hypothesis testing, multiple testing correction, analysis code execution) only supports DESeq2.

[b] TSPM uses a two-stage Poisson framework, applying a quasi-likelihood adjustment when overdispersion is detected.

[c] BH can be applied explicitly to the raw p-values from all listed methods; BH correction is applied separately within each primary contrast, with significance defined as an adjusted $p$-value $< 0.05$.

#### Section 1.3 statLens Realworld Benchmark

**Supplementary Table 1.3** Distribution of the 102 MCQ evaluation tasks across five omics feature types.

| Omics category | Representative modalities or assay types | N datasets | N tasks | % of tasks |
|---|---|---|---|---|
| Microbiome | 16S rRNA gene amplicon sequencing (V3–V4 or V4 region), profiled as DADA2 amplicon sequence variants (ASVs) or 97% OTUs | 3 | 7 | 6.86% |
| Proteomics | Label-free data-dependent LC-MS/MS (timsTOF dda-PASEF, Orbitrap Fusion) and aptamer-based affinity profiling (SomaScan v4) | 3 | 7 | 6.86% |
| Metabolomics | Untargeted and targeted LC-MS/MS (reversed-phase and HILIC; MRM/SRM), GC-MS, and $^{1}$H-NMR metabolite profiling, and lipidomics (LC-MS and direct-infusion shotgun MS/MS) | 14 | 24 | 23.53% |
| Transcriptomics | Bulk RNA sequencing (poly(A)/mRNA-seq, rRNA-depleted total RNA-seq, and 3′ RNA-seq) provided as raw gene-level read or UMI counts | 49 | 57 | 55.88% |
| Other | DNA methylation (targeted bisulfite sequencing and the Illumina HorvathMammalMethylChip40 array) and small RNA profiling (small RNA sequencing and the Affymetrix GeneChip miRNA 4.0 array) | 5 | 7 | 6.86% |
| Total | – | 74 | 102 | 100.00% |

**Supplementary Table 1.4** Distribution of the 26 bulk RNA-seq tasks with a DESeq2 gold standard across source organisms.

| Organism | Representative tissues or cell types | N datasets | N tasks | % of tasks |
|---|---|---|---|---|
| *Homo sapiens* | Cancer and immortalised cell lines (small cell lung cancer, leukaemia, colorectal cancer, iSLK, and HaCaT keratinocytes), embryonic stem cells, stem cell-derived neurons and thymic epithelial progenitors, astrocytes, endometrial stromal cells, PBMCs, and lung tissue | 15 | 18 | 69.23% |
| *Mus musculus* | MIN6 insulinoma cells, kidney, and splenic $CD4^{+}$ T cells | 3 | 3 | 11.54% |
| Other | *Candida albicans* and bacteria (*Arthrobacter* sp., *Chlamydia trachomatis*, and *Vibrio cholerae*) | 4 | 5 | 19.23% |
| Total | – | 22 | 26 | 100.00% |

### Section 2. statLens design card

**Supplementary Table 2.1** Fields in the `statLens` Design Card (part 1 of 3).

| Field | Type | Description |
|---|---|---|
| *2.0 Data Description* | | |
| `Matrix samples` | int | Number of samples (columns) in the omics feature matrix. |
| `Matrix genes` | int | Number of genes (rows) in the omics feature matrix. |
| `Matrix gene ID column` | text | Name of the matrix column that holds the gene IDs. |
| `Omics data type` | text | Kind of values in the matrix (whole or continuous numbers, with or without negatives) and their range. |
| `Metadata` | text | Number of rows and columns in the metadata table. |
| `Matrix samples found in Sample ID column` | text | Number of matrix samples found in the sample ID column of the metadata. |
| `Samples per group level` | text | Number of samples at each level of the main condition variable. |
| `Samples per time value` | text | Number of samples at each value of the time variable. |
| `Units` | text | Number of biological units and samples per unit, from the subject ID column. |
| `Samples kept by the subset` | text | Number of samples kept by the subset definition. |
| *2.1 Experimental background and variable description* | | |
| `Biological system` | text | Tissue, cell type, disease, or treatment background of the study. |
| `Biological unit` | text | What one sample represents, e.g., one blood draw from one patient. |
| `Sample ID column` | text | Metadata column that holds the sample IDs of the matrix. |
| `Main condition variable` | text | Metadata column of the main group or treatment, if any. |
| `Group label meanings` | dict | Meaning of each level of the main condition variable. |
| `Time variable` | text | Metadata column of the sampling time, if any. |
| `Time meaning and unit` | text | Meaning and unit of the time values. |

**Supplementary Table 2.1 (continued).** Fields in the `statLens` Design Card (part 2 of 3).

| Field | Type | Description |
|---|---|---|
| *2.2 Research question and comparison setup* | | |
| `Research question` | text | Scientific question the analysis is to answer. |
| `What the question compares` | enum | A difference between groups, a change over time, how the change over time differs between groups, or something else (out of scope). |
| `Comparison group` | text | Group compared with the reference group, when two specific groups are compared. |
| `Reference group` | text | Group against which the comparison group is measured. |
| `Analysis sample range` | enum | All samples, or a specified subset. |
| `Subset definition` | dict | Metadata column and values that define the subset. |
| *2.3 Sample dependence and technical replicates* | | |
| `Shared biological origin` | enum | Whether several samples come from the same biological unit (yes or no). |
| `Subject / biological unit ID column` | text | Metadata column that links the samples of one biological unit. |
| `Sampling relationship` | text | How the samples of one biological unit relate to one another, e.g., repeated blood draws. |
| `Technical replicate status` | enum | Whether technical replicates are absent, already merged, or not merged yet. |
| *2.4 Batch and other variables* | | |
| `Batch columns` | list | Metadata columns that record a technical batch. |
| `Batch meanings` | dict | Technical process that each batch column stands for. |
| `Relevant variable columns` | list | Other metadata columns that may matter, e.g., age or sex. |
| `Variable meanings` | dict | Meaning and type (categorical or continuous) of each relevant variable. |
| `Pre-specified adjustment requirements` | text | Adjustments fixed in advance by the study; optional. |

**Supplementary Table 2.1 (continued).** Fields in the `statLens` Design Card (part 3 of 3).

| Field | Type | Description |
|---|---|---|
| *2.5 Data context* | | |
| `Measurement technology` | enum | Technology that produced the matrix, e.g., bulk RNA-seq, single-cell RNA-seq, or microarray. |
| `Feature type` | enum | What one row of the matrix is: gene, transcript, or other count-based feature. |
| `Value unit` | enum | What one matrix value is before any log transformation, e.g., raw counts, TPM, or intensity. |
| `Log scale` | enum | Whether the matrix values are linear or log-transformed. |
| `Other processing` | text | Any other processing of the values, e.g., pseudobulk aggregation, filtering, or batch correction; optional. |
| *2.6 Primary contrast* | | |
| `Primary contrast` | enum | Primary contrast derived by the code: group-by-time (GT), time course (TC), multi-group (MG), basic two-group (BG), or out of scope (OOS). |
| `Group column` | text | Group column used by the primary contrast. |
| `Time column` | text | Time column used by the primary contrast. |
| `Comparison` | text | Statement of the comparison, with the actual column names and levels. |
| `Null hypothesis` | text | Null hypothesis of the primary contrast, in formal and plain-language form. |
| `Alternative hypothesis` | text | Alternative hypothesis of the primary contrast, in formal and plain-language form. |
| *2.7 Subject dependence* | | |
| `Subject dependence` | enum | Subject dependence derived by the code: independent (IND), paired or repeated (PR), or out of scope (OOS). |
| `Subject column` | text | Metadata column that identifies the subjects, for paired or repeated designs. |
| `Description` | text | How the samples depend on one another through shared subjects. |
| `Subject term` | text | Subject term of the model (none, or subject fixed effects), in formal and plain-language form. |

*Note:* Each row also records its provenance: entered by the user, filled in by the model, derived by the code, or unknown. The Data Description is counted from the two input files; the primary contrast and the subject dependence are derived by the code from the answers and the metadata, and the user can change them. Fields that do not apply are left out. The card does not contain a statistical package, model family, or test.

### Section 3. statLens model formulation

This section describes how the five `statLens` design axes (statistical model family, primary comparison, subject dependence, batch effects and other covariates) are represented as model components and inferential targets. The formulation covers the negative-binomial GLMs, Gaussian linear models, Poisson GLMs and beta regression considered by the Reasoning module. These models share a representation of the conditional mean through a linear predictor, with family-specific distributions and link functions. The current public release implements model fitting and hypothesis testing through DESeq2 for negative-binomial models.

#### 3.1 Shared mean structure

Let $Y_{ij}$ denote the response for molecular feature $i$ in analysed sample $j$, on the selected modelling scale, and let $\mu_{ij}$ denote its conditional mean. The linear predictor is

$$\begin{aligned}\eta_{ij} = h_f(\mu_{ij}) = o_{ij} + \beta_{0i} + \mathcal{P}_i(g_j, t_j) \\ + I_S \mathbf{s}_j^\top \boldsymbol{a}_i + I_B \mathbf{b}_j^\top \boldsymbol{\theta}_i + I_C \mathbf{c}_j^\top \boldsymbol{\gamma}_i,\end{aligned} \tag{1}$$

where $h_f$ is the link function for family $f$, $\beta_{0i}$ is a feature-specific intercept, and $\mathcal{P}_i(g_j, t_j)$ represents the primary-design terms involving group $g_j$, time $t_j$, or their interaction. The vectors $\mathbf{s}_j$, $\mathbf{b}_j$ and $\mathbf{c}_j$ encode subject membership, batch membership and other covariates, respectively; $\boldsymbol{a}_i$, $\boldsymbol{\theta}_i$ and $\boldsymbol{\gamma}_i$ are their feature-specific coefficients. The indicators $I_S, I_B, I_C \in \{0, 1\}$ record whether these adjustment terms are included in the confirmed specification.

The term $o_{ij}$ is a method-specific offset and is zero when no offset is used. Its definition depends on the measurement and normalization procedure. An offset may be estimated during data preparation or analysis, but is treated as a supplied adjustment within the mean-model regression. It is excluded from the regression-coefficient vector and is not directly tested. For the DESeq2 implementation, the offset is determined by the estimated size factors, as described below.

Equation 1 identifies the conceptual model components. Their separately identifiable representation depends on the observed design and variable coding.

#### 3.2 Observation models

Conditional on the included design variables, the candidate observation models are

$$Y_{ij} \sim \begin{cases} \mathrm{NB}(\mu_{ij}, \alpha_i), & \text{negative-binomial GLM,} \\ \mathcal{N}(\mu_{ij}, \sigma_i^2), & \text{Gaussian linear model,} \\ \mathrm{Poisson}(\mu_{ij}), & \text{Poisson GLM,} \\ \mathrm{Beta}(\mu_{ij}\phi_i, (1-\mu_{ij})\phi_i), & \text{beta regression.} \end{cases} \tag{2}$$

The negative-binomial and Poisson models use a log link, the Gaussian model uses an identity link, and the beta mean model uses a logit link. Here, log denotes the natural logarithm. The negative-binomial dispersion $\alpha_i > 0$ gives

$$\mathrm{Var}(Y_{ij} \mid \text{design}) = \mu_{ij} + \alpha_i \mu_{ij}^2,$$

and $\sigma_i^2 > 0$ is the Gaussian residual variance. For beta regression, $\phi_i > 0$ is a feature-specific precision parameter, giving conditional variance $\mu_{ij}(1-\mu_{ij})/(1+\phi_i)$.

Count models apply to responses on the count scale. Gaussian models apply on the confirmed analysis scale, including any specified transformation. Ordinary beta regression requires responses strictly within $(0, 1)$. Model-family suitability therefore depends on the measurement process and preprocessing as well as numerical properties. The dispersion, residual-variance and precision parameters are nuisance parameters for the mean comparisons considered here.

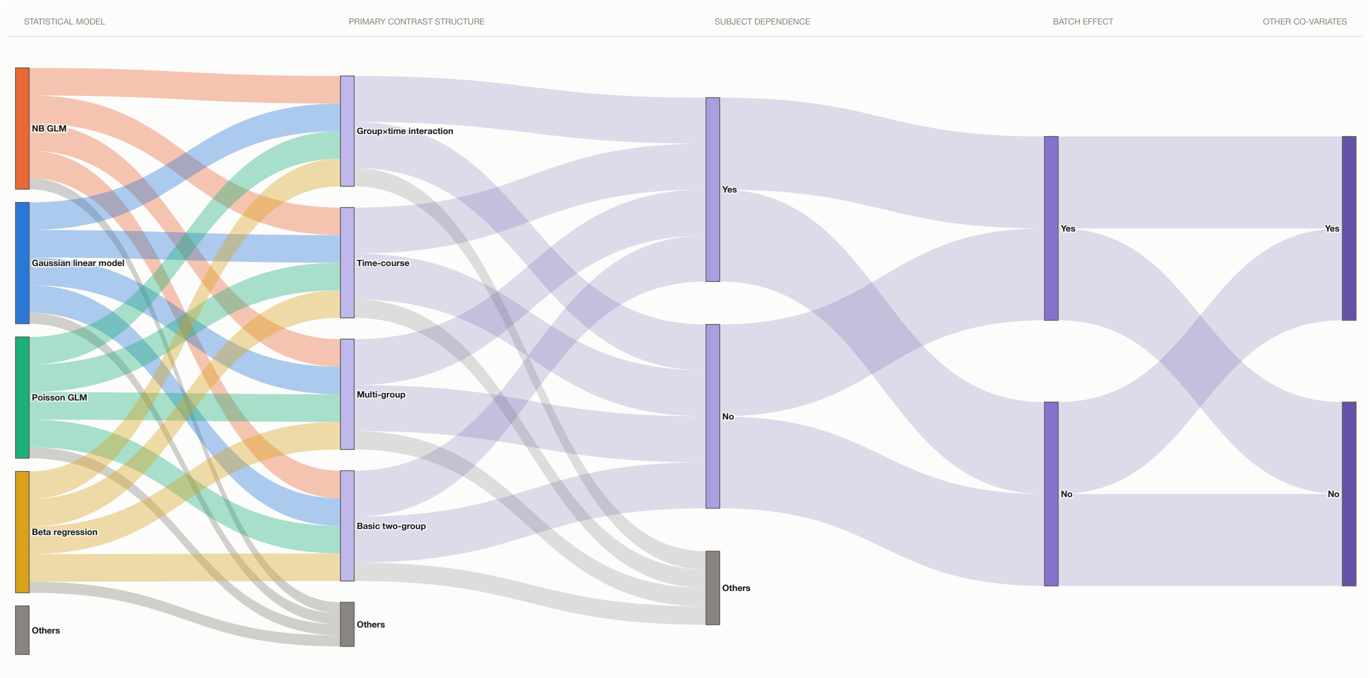


**Supplementary Figure 3.1. Compositional organization of the supported differential expression analysis** The schematic combines statistical model, primary DEA contrast structure, subject dependence, batch effect states, and other-covariate states to define supported workflow configurations.

#### 3.3 Primary comparison and adjustment terms

For basic two-group comparisons (BG) and multi-group comparisons (MG), the primary-design component includes the relevant group terms. Time-course designs (TC) include the confirmed time representation. Group-by-time designs (GT) include group and time terms and their interaction, subject to an identifiable parameterization. Categorical time encodes separate time-point effects, whereas numeric time specifies a trend on the linear-predictor scale. The time encoding must therefore agree with the intended question.

The primary-design class does not uniquely determine the inferential target. For example, a multi-group model can support a specified pairwise comparison or a joint test of group differences. The target comparison is defined separately from reference-level coding and is expressed in the executable coefficient parameterization.

Subject dependence is represented through subject fixed effects in supported paired or repeated designs. These terms adjust for subject-specific baseline differences; the formulation does not specify an additional within-subject covariance model. Batch effects and other covariates enter as confirmed adjustment terms. Their inclusion and coding must be supported by the analysed sample structure.

#### 3.4 Design validation and executable parameterization

The reviewed input formula from step 4 defines a candidate design matrix $X_{\text{in}}$. In the current execution path, step 6 constructs and validates an executable matrix $X$ using the metadata for the analysed samples. For feature $i$, the executable representation is

$$\boldsymbol{\eta}_i = \boldsymbol{o}_i + X\boldsymbol{\beta}_i,$$

where each row of $X$ corresponds to one analysed sample, and $\boldsymbol{\beta}_i$ contains the mean-model coefficients in the validated parameterization.

When input terms are redundant, an equivalent executable parameterization may remove redundant representations or introduce explicit derived columns. For the same analysed samples and offsets, equivalence requires

$$\mathrm{col}(X_{\mathrm{in}}) = \mathrm{col}(X),$$

where $\mathrm{col}(\cdot)$ denotes column space. The intended comparison must also be mapped consistently into the executable coding. These conditions preserve the modelled mean structure and the estimable target.

For example, when group membership is constant within subjects, group main effects can be redundant with subject fixed effects. Group-by-time comparisons may nevertheless remain estimable when supported by the observed measurements. An executable parameterization can absorb the redundant group representation into the subject terms while retaining the relevant interaction columns.

Validation assesses matrix rank, residual degrees of freedom and estimability of the primary comparison. A contrast in the input coefficient parameterization is estimable only if its coefficient vector lies in the row space of $X_{\mathrm{in}}$. Reparameterization preserves estimable functions but cannot identify a comparison unsupported by the data. Residual degrees of freedom are

$$d_{\mathrm{res}} = n - \mathrm{rank}(X),$$

where $n$ is the number of analysed samples. This quantity describes the remaining design dimension and is distinct from the degrees of freedom of a hypothesis test.

#### 3.5 Hypotheses and DESeq2 implementation

For an estimable primary comparison, the feature-specific hypotheses are

$$H_{0i} : \mathbf{L}^\top \boldsymbol{\beta}_i = \mathbf{0}, \qquad H_{1i} : \mathbf{L}^\top \boldsymbol{\beta}_i \neq \mathbf{0}.$$

Here, $\mathbf{L} \in \mathbb{R}^{p\times r}$ specifies $r$ linearly independent restrictions on the $p$ executable mean-model coefficients. A contrast vector defines a single comparison; a contrast matrix defines a joint hypothesis. The alternative means that at least one tested component differs from zero. The scientific target is retained across equivalent coding choices by updating $\mathbf{L}$ accordingly.

For the public DESeq2 execution path, the negative-binomial model is written as

$$Y_{ij} \sim \mathrm{NB}(\mu_{ij}, \alpha_i), \qquad \mu_{ij} = s_j q_{ij}, \qquad \log_2(q_{ij}) = \mathbf{x}_j^\top \boldsymbol{\beta}_i^{(2)},$$

where $s_j$ is the sample-specific size factor and $\mathbf{x}_j^\top$ is row $j$ of $X$. The coefficients $\boldsymbol{\beta}_i^{(2)}$ are expressed on the log-base-2 scale; the natural-log coefficients satisfy $\boldsymbol{\beta}_i = (\log 2)\boldsymbol{\beta}_i^{(2)}$. The corresponding offset is $\log(s_j)$. Size factors are estimated using the configured method and are not generally equal to raw sample totals[?] .

A Wald test evaluates a specified contrast in the executable coding. An LRT compares nested full and reduced models fitted to the same samples, with the reduced model representing the intended null hypothesis. For an equivalent set of $r$ independent constraints, the LRT degrees of freedom are

$$d_{\mathrm{LRT}} = \mathrm{rank}(X_{\mathrm{full}}) - \mathrm{rank}(X_{\mathrm{reduced}}) = r.$$

The choice of contrast or reduced model determines which effect is tested. Numerical model fitting and hypothesis testing are performed when the analysis script is executed in step 7.

### Section 4. statLens evaluation

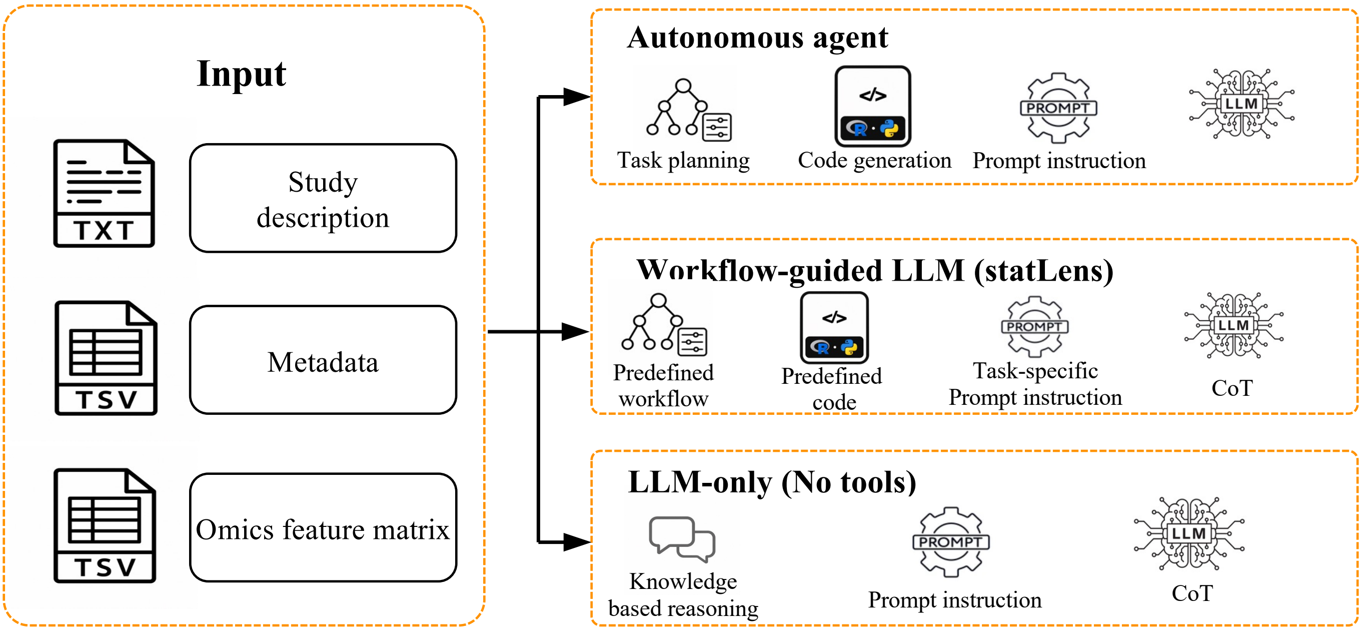


**Supplementary Figure 4.1. statLens evaluation pipeline**

| Answer | Reasoning | Correctness |
|---|---|---|
| **DATA MODALITY** *What do the values in omics_feature_matrix.tsv represent, as they are provided?* | | |
| **B. Non-normalized read or sequence counts** | The matrix holds whole-number, non-normalized read-pair counts per gene, so the values are raw counts." *All values are integers from 0 to 2,663,883, and the study description says featureCounts counts were rounded after fractional multi-mapping.* | Correct |
| **STAITSTICAL MODEL** *Which contrast structure matches the primary comparison?* | | |
| **D. Negative binomial GLM on counts** | Over-dispersed low-input RNA-seq counts are best tested gene by gene with a negative binomial GLM." *A Gaussian model after a log-type transformation (voom/limma) is also acceptable; Poisson is not, because it assumes variance equals the mean.* | Correct |
| **PRIMARY CONTRAST** *Which structure best describes the primary question of this study, as the study description states it?* | | |
| **D. Basic two-group comparison (BG)** | The study asks only whether aged_Ri differs from aged_control, which is a two-group comparison even though the study has three conditions." *young_control is explicitly outside the primary question.* | Correct |
| **SAMPLE DEPENDENCE** *Should a batch effect be adjusted?* | | |
| **B. Treat the analysed samples as independent: no subject or unit term (IND)** | Each sample is an independent culture with its own library and no recorded subject link, so no unit term belongs in the model." *The numbers 1 to 3 in the sample names only count samples within each condition, so they do not identify pairs.* | Correct |
| **FORMULA** *Which model formula, together with the hypothesis it tests, should be used for the primary question?* | | |
| **B. ~ condition** | With independent samples and no informative covariate, the model needs only the condition term to compare aged_Ri with aged_control." *Strain is C57BL/6 for every sample, and the sequencing-batch codes are unexplained, so neither is a valid covariate.* | Correct |

**Supplementary Figure 4.2. statLens evaluation example**

Table 4.1 MCQ accuracy of autonomous agent approach.

| Model | Overall | RNA-seq | Metabolomics | Proteomics | Microbiome | Others |
|---|---|---|---|---|---|---|
| GPT-5.5 | 99.5 (99.6, 99.4) | 99.9 (100.0, 99.6) | 100.0 (100.0, 100.0) | 99.0 (100.0, 97.1) | 95.2 (97.1, 94.3) | 100.0 (100.0, 100.0) |
| DeepSeek-V4.1-Flash | 98.9 (99.2, 98.4) | 99.6 (99.6, 99.6) | 98.9 (100.0, 98.3) | 100.0 (100.0, 100.0) | 92.4 (94.3, 88.6) | 98.1 (100.0, 97.1) |
| Claude Haiku 5.5 | 97.5 (97.8, 97.3) | 99.6 (99.6, 99.6) | 94.4 (95.0, 94.2) | 95.2 (100.0, 91.4) | 95.2 (97.1, 94.3) | 95.2 (97.1, 91.4) |
| Gemma 4 26B-A4B | 92.2 (93.1, 91.2) | 98.0 (98.6, 97.2) | 84.4 (86.7, 82.5) | 81.0 (82.9, 80.0) | 84.8 (88.6, 82.9) | 89.5 (91.4, 88.6) |
| Qwen3.5-9B | 89.2 (90.0, 88.8) | 95.6 (96.8, 94.4) | 81.7 (85.0, 79.2) | 75.2 (82.9, 62.9) | 86.7 (88.6, 85.7) | 80.0 (82.9, 77.1) |
| Phi-4 | 84.1 (86.5, 82.2) | 90.8 (92.3, 89.5) | 76.7 (78.3, 74.2) | 71.4 (80.0, 60.0) | 79.0 (82.9, 71.4) | 72.4 (77.1, 68.6) |
| Gemma 3 12B | 58.7 (59.6, 57.8) | 67.5 (69.1, 66.7) | 53.6 (62.5, 43.3) | 46.7 (57.1, 34.3) | 41.0 (48.6, 31.4) | 34.3 (51.4, 20.0) |

Table 4.2 MCQ accuracy of LLM-only approach.

| Model | Overall | RNA-seq | Metabolomics | Proteomics | Microbiome | Others |
|---|---|---|---|---|---|---|
| GPT-5.5 | 36.1 (37.1, 34.7) | 39.6 (42.8, 36.5) | 30.6 (35.8, 23.3) | 39.0 (40.0, 37.1) | 28.6 (34.3, 22.9) | 31.4 (34.3, 28.6) |
| DeepSeek-V4.1-Flash | 46.8 (47.6, 46.1) | 54.9 (56.8, 53.7) | 31.4 (32.5, 30.0) | 34.3 (40.0, 31.4) | 48.6 (48.6, 48.6) | 44.8 (51.4, 40.0) |
| Claude Haiku 5.5 | 42.4 (42.9, 42.0) | 43.3 (43.5, 43.2) | 41.1 (41.7, 40.8) | 44.8 (51.4, 40.0) | 35.2 (40.0, 28.6) | 43.8 (48.6, 40.0) |
| Gemma 4 26B-A4B | 17.5 (18.2, 17.1) | 17.9 (18.2, 17.2) | 20.0 (20.8, 19.2) | 15.2 (20.0, 11.4) | 7.6 (8.6, 5.7) | 18.1 (22.9, 14.3) |
| Qwen3.5-9B | 43.5 (45.1, 42.2) | 54.4 (55.4, 53.7) | 21.9 (22.5, 20.8) | 29.5 (37.1, 25.7) | 41.9 (45.7, 37.1) | 43.8 (48.6, 40.0) |
| Phi-4 | 39.2 (39.8, 38.2) | 47.6 (48.8, 46.3) | 21.7 (24.2, 20.0) | 33.3 (37.1, 31.4) | 37.1 (40.0, 34.3) | 38.1 (40.0, 34.3) |
| Gemma 3 12B | 40.1 (41.0, 39.2) | 42.3 (44.6, 40.7) | 37.2 (38.3, 36.7) | 40.0 (45.7, 37.1) | 30.5 (34.3, 28.6) | 41.9 (42.9, 40.0) |

Table 4.3 MCQ accuracy of statLens approach.

| Model | Overall | RNA-seq | Metabolomics | Proteomics | Microbiome | Others |
|---|---|---|---|---|---|---|
| GPT-5.5 | 95.0 (96.7, 93.9) | 98.2 (99.3, 97.5) | 93.6 (97.5, 91.7) | 81.0 (82.9, 80.0) | 100.0 (100.0, 100.0) | 81.9 (82.9, 80.0) |
| DeepSeek-V4.1-Flash | 94.1 (95.1, 93.3) | 97.2 (98.2, 96.1) | 92.8 (93.3, 92.5) | 81.0 (82.9, 80.0) | 98.1 (100.0, 97.1) | 82.9 (82.9, 82.9) |
| Claude Haiku 5.5 | 94.8 (97.5, 93.1) | 97.5 (99.6, 95.8) | 93.9 (97.5, 91.7) | 81.9 (82.9, 80.0) | 98.1 (100.0, 97.1) | 84.8 (91.4, 80.0) |
| Gemma 4 26B-A4B | 93.9 (95.3, 92.9) | 97.4 (98.6, 96.1) | 94.7 (97.5, 93.3) | 81.9 (82.9, 80.0) | 88.6 (88.6, 88.6) | 80.0 (80.0, 80.0) |
| Qwen3.5-9B | 94.4 (96.3, 93.1) | 97.0 (98.6, 95.4) | 94.4 (97.5, 92.5) | 81.0 (82.9, 80.0) | 98.1 (100.0, 97.1) | 82.9 (82.9, 82.9) |
| Phi-4 | 93.7 (95.5, 92.7) | 97.7 (99.3, 96.5) | 93.9 (97.5, 91.7) | 76.2 (77.1, 74.3) | 88.6 (88.6, 88.6) | 82.9 (85.7, 80.0) |
| Gemma 3 12B | 92.5 (93.7, 91.8) | 96.4 (97.2, 95.4) | 91.4 (94.2, 90.0) | 81.0 (82.9, 77.1) | 88.6 (88.6, 88.6) | 80.0 (80.0, 80.0) |

### Section 5. statLens software

#### Section 5.1 statLens software workflow

| Step | Module | Core function | Responsibility |
|---|---|---|---|
| **1 Information collection** | Perception | Profile the matrix and extract study information | Deterministic data profiling; evidence-grounded LLM extraction with programmatic verification |
| **2 Design Card** | Perception | Define data context, research question, and study design | Rule-based derivation and constrained LLM proposals, with user-confirmed decisions |
| **3 Statistical model recommendation** | Reasoning | Recommend the statistical model family | Fact-grounded LLM assessment, programmatic consistency checks, and final user selection |
| **4 Model specification** | Reasoning | Generate and validate the design formula | Deterministic formula generation and validation, with user-editable specifications |
| **5 statLens Insight** | Reasoning | Explain decisions and limitations | LLM-generated explanations with code check for structural and coverage |
| **6 Model fitting** | Execution | Select statistical test and analysis parameters | Rule-based test recommendations and parameter validation, with user-defined settings |
| **7 Analysis code** | Execution | Generate the R script (current version only support **DESeq2**) | Template-based code generation, with user-editable scripts and user-initiated execution |

**Supplementary Table 5.1. statLens software workflow demonstration**

#### Section 5.2 statLens output package

After the finalized script is executed, statLens returns a standardized result package containing:

1. `results.csv`, the complete differential-analysis table, including feature identifiers, $\log_2$ fold changes, standard errors, test statistics, nominal $P$ values, and adjusted $P$ values;
2. `significant_features.csv`, a filtered table of significant molecular features based on the workflow-specific false-discovery-rate threshold;
3. `volcano_plot.png`, displaying $\log_2$ fold change against $-\log_{10}$ adjusted $P$ value;
4. `MA_plot.png`, displaying $\log_2$ fold change against mean expression or abundance to assess abundance-dependent effects;
5. `PCA_plot.png`, summarizing sample-level variation in the normalized expression or abundance matrix;
6. `top_DE_features_heatmap.png`, displaying the top differentially expressed or abundant features across samples; and
7. `topK_DE_features_barplot.png` (default $K = 20$), displaying the top features ranked by absolute $\log_2$ fold change.

#### Section 5.3 statLens Portal

To support community adoption and continued development of statLens, we have established the `statLens` Portal, available at https://statlens.ai. The portal comprises two core components. The first, *Community Methods*, enables and encourages researchers to contribute advanced statistical methodologies along with their source code, example input and output data, documentation, code repositories, and associated citations. By curating these contributions in a standardized, discoverable format, the component aims to accelerate the translation of methodological research into practical application across the research community while ensuring contributing authors receive appropriate attribution and citation whenever their methods are used. The second component, *Run statLens Agent*, provides an agentic interface that assists users in selecting appropriate statistical methods and models for their study design and data, and allows them to execute the recommended analyses directly on the platform. Contributed methods remain credited to their original developers; any fees associated with running analyses support platform maintenance and the computational resources required for execution, not the contributed methods themselves. The portal is under active development, and additional components and data types will be supported in future releases.
